# Defense-Suppressive Fragments of RIN4 generated by AvrRpt2 Participate in NDR1-dependent Activation of RPS2

**DOI:** 10.1101/2025.06.02.656948

**Authors:** Ahmed J. Afzal, Maheen Alam, Jianhua Huang, Moneeza Akbar Agha, Luis da Cunha, Muneeza Iqbal Rai, Jibran Tahir, Anmbreen Jamroze, Jonathan D. G. Jones, David Mackey

## Abstract

Plant nucleotide-binding, leucine-rich-repeat (NLR) immune receptors recognize pathogen effectors and activate immunity. The NLR RPS2 recognizes AvrRpt2, a *Pseudomonas* effector that promotes virulence by proteolytically cleaving a membrane-tethered host protein, RIN4. RIN4 cleavage by AvrRpt2 generates fragments that activate RPS2. A model for RPS2 activation by RIN4 destruction is consistent with the ectopic activity of RPS2 in plants lacking RIN4 but does not explain the link between AvrRpt2’s virulence activity and RPS2 activation. We found that non-membrane-tethered RIN4 derivatives are potent cytosolic activators of RPS2. Activation of RPS2 by these RIN4 derivatives, like AvrRpt2-induced activation, and unlike ectopic activation in the absence of RIN4, requires the defense signaling protein NDR1. Cleavage products of RIN4 produced by AvrRpt2 play contrasting roles in the activation of RPS2, with the membrane-tethered C-terminal fragment suppressing RPS2 and the non-membrane-tethered internal fragment, dependent on compatibility with the C-terminal fragment, overcoming its suppression of RPS2.

**Highlights:**

- **Non-membrane tethered derivatives of RIN4 activate RPS2-induced cell death**
- **Activation of RPS2 by non-membrane-tethered derivatives of RIN4 requires NDR1**
- **AvrRpt2-induced cleavage fragments of RIN4 play contrasting roles in RPS2 activation**

## Introduction

Plants are attacked by diverse pathogens that must overcome plant innate immunity to cause disease. Activation of immunity relies on receptors that detect pathogen-related molecules either inside or outside the plant cell. Plants utilize cell surface pattern recognition receptors (PRRs) to activate Pattern-Triggered Immunity (PTI) upon recognition of conserved microbial features known as Microbe or Pathogen Associated Molecular Patterns (MAMPs or PAMPs) ^1–5^. Gram-negative bacteria utilize a specialized needle-like type three secretion system (TTSS) to deliver effector proteins into host cells to promote effector-triggered susceptibility (ETS) by suppressing host immunity, *e.g.*, PTI, and promoting a nutritious water-rich apoplast environment ^3,6–10^. Effectors can be detected by plant NLRs which activates a rapid immune response called effector-triggered immunity (ETI) that often activates localized cell death called the hypersensitive response (HR) ^3,10,11^. In recent years, it has been demonstrated that PTI and ETI work synergistically to strengthen plant defenses against pathogen infection ^12,13^. Activation of both systems together provides more effective resistance, highlighting the importance of mutual potentiation of cell surface and intracellular receptors for an effective plant defense system ^12,13^.

NLRs contain central nucleotide binding and C-terminal leucine rich repeat domains and typically N-terminal a coiled coil (CNLs) ^14^ or toll-interleukin receptor (TNLs) domains ^15–18^. Activation of an NLR by its corresponding effector can occur either through a direct interaction, sometimes with an integrated domain acting as a decoy of the effector’s actual virulence target, or indirectly through an effector-induced modifications of a *bona fide* host target or a decoy ^3,19–21^. In the latter case, the NLR is said to “guard” the host target by monitoring for its perturbation by an effector protein ^22^. A particularly well-studied example of a *bona fide* virulence target that is guarded by multiple NLRs is RPM1-interacting protein 4 (RIN4).

RIN4 is a 211 amino acid, intrinsically unstructured protein that contains two Nitrate Induced (NOI) domains of unknown function and is anchored to the plasma membrane by lipidation of C-terminal cysteine residues ^23–26^. RIN4 negatively regulates PTI, with overexpression attenuating PTI in Arabidopsis plants ^24,27,28^. Multiple *P. syringae* T3Es, including AvrRpm1, AvrB, AvrRpt2, HopF2 and HopZ3, target and modify RIN4 to compromise host defenses and promote bacterial growth ^29–33^. For example, AvrB induces RIN4-interacting protein kinase (RIPK) ^34,35^ mediated phosphorylation of RIN4 at T166 within the C-terminal NOI (C-NOI) domain; which counteracts activation of PTI associated with MAMP-induced phosphorylation of another key residue of RIN4, S141, also within the C-NOI ^36^. Another example is the proteolytic cleavage of RIN4 by AvrRpt2 within each of the two NOI domains, which generates three fragments (AvrRpt2-cleavage products, ACP1-3) ^23,37,38^. Both the soluble ACP2 and membrane-tethered ACP3, which contain truncated N-NOI and C-NOI domains, respectively, persist following their generation by AvrRpt2 and are hyperactive suppressors of PTI relative to intact RIN4 ^27^. Thus, it is well established that effector-induced perturbations of RIN4, including the NOI domains, contribute to suppression of plant defense.

In addition to regulating PTI, RIN4 also regulates ETI via interaction with and effector-induced activation of multiple, evolutionarily unrelated NLRs ^24,29,30,35,36,39–43^. A central prediction of the guard hypothesis is that the virulence-associated modification of a guardee will trigger the activation of an associated NLR ^22^. Consistent with this prediction, modifications of RIN4 induced by AvrRpm1 or AvrB activate a plasma membrane localized CNL, Resistance to *Pseudomonas syringae* pv. *maculicola* 1 (RPM1) ^35^. In the absence of effector activation, RIN4 interacts with and negatively regulates RPM1 at the plasma membrane ^29,44–46^. Through perturbation of RIN4, AvrB and AvrRpm1 activate the soybean CNLs Rpg1b and Rpg1r, respectively, and AvrRpm1 also weakly activates another Arabidopsis CNL, Resistance to *Pseudomonas syringae*2 (RPS2) ^47^. RPS2, an apple CNL MR5, and a tomato CNL Ptr1, each respond to proteolysis of RIN4 by homologs of AvrRpt2 ^30,42,43^. Remarkably, *RPM1*, *Rpg1b*, *Rpg1r*, *RPS2*, *MR5,* and *Ptr1* each evolved independently ^40,42,43^. Collectively, these findings highlight the importance of “guarding” RIN4 for plant immunity.

The mechanism underlying how virulence-associated proteolysis of RIN4 by AvrRpt2 activates RPS2 remains obscure. In Arabidopsis plants lacking RIN4, RPS2 is ectopically active, *i.e.* active in the absence of an activating T3E ^30,48^. Thus, a simple interpretation is that elimination of RIN4 by AvrRpt2 activates RPS2 ^30,48,49^. However, unlike ectopic activation of RPS2, activation of RPS2 by AvrRpt2 requires the RIN4-interacting protein NDR1 (NON-RACE-SPECIFIC DISEASE RESISTANCE1) ^46,50–52^. MR5 is not ectopically active and is activated by the ACP3 fragment of apple RIN4 ^42^. The ACP2 and ACP3 fragments, which are defense-suppressive in Arabidopsis, are candidates for involvement in the NDR1-dependent activation of RPS2.

Activated CNLs, such as ZAR1, can induce cell death by oligomerizing into a multimeric complex known as a resistosome that disrupts plasma membrane integrity ^53–56^. Resistosome formation requires the conserved MADA motif present in the N-terminus α1 helix ^57,58^. RPS2 lacks a canonical MADA motif and, like other MADA-less CNLs, relies on ‘helper’ NLRs to carry out the immune response ^57,59,60^. The *RESISTANCE TO POWDERY MILDEW 8* (*RPW8*) or *Activated Disease Resistance 1* (*ADR1*) genes encode such helpers ^60–62^. ETI upon activation of RPS2 by AvrRpt2 is significantly attenuated in *adr1* triple mutant plants indicating that the ADR1 helpers act downstream RPS2 ^60^. We speculate that, upon sensing the AvrRpt2-induced perturbation of RIN4, activated RPS2 propagates the defense signal by interacting with ADR1.

We observed previously that RIN4 derivatives not tethered to the plasma membrane, because they lack the C-terminal fatty acylation motif, are more potent suppressors of PTI and also elicit an HR-like cell death response ^27^. We speculated that these non-membrane-tethered derivatives of RIN4 might elicit an HR by functioning in a manner analogous to the soluble ACP2 fragment produced by AvrRpt2. Here we show that non-membrane-tethered derivatives of RIN4, which localize to both the cytosol and nucleus, function in the cytosol to activate RPS2. Similar to AvrRpt2-dependent activation, RPS2 activation by non-membrane-tethered derivatives of RIN4 requires NDR1. ACP2 and ACP3 play contrasting roles in RPS2 regulation. Membrane-tethered ACP3 retains the ability to inhibit ectopic activation of RPS2. Conversely, non-membrane-tethered ACP2, which localizes to both the cytosol and nucleus, functions outside the nucleus to activate RPS2 by overcoming suppression mediated by ACP3. Compatibility between ACP2 and ACP3 is required for ACP2 to overcome suppression by ACP3, indicating that ACP2 likely interacts with a pre-activation complex containing RPS2 and ACP3. Consistent with predictions based on the “guard hypothesis”, these findings link the virulence-associated perturbation of RIN4 by AvrRpt2 to the activation of RPS2.

## Results

### Cell death caused by non-membrane-tethered derivatives of RIN4 is dependent on RPS2

RIN4 contains a C-terminal fatty acylation site that includes three cysteines at residues 203-205 ^23^. Palmitoylation of one or more of these C-terminal cysteines causes plasma membrane localization of RIN4 ^23,24^. Three non-membrane-tethered derivatives of RIN4 (177Δ211, 203Δ211, and CCC>AAA) that lack the C-terminal acylation site **(Figure 1A)** elicit a macroscopic cell death response when expressed as transgenes under control of a dexamethasone (dex)-inducible promoter in the *Arabidopsis* ecotype Col-0 **(Figure S1A-C)**. The strong cell death response elicited by non-membrane-tethered RIN4 derivatives cannot be attributed to expression levels; anti-T7 immunoblotting demonstrated that RIN4FL, which induces only modest chlorotic symptoms, accumulates to significantly higher levels than the non-membrane-tethered RIN4 derivatives ^27^.

**Figure 1.**
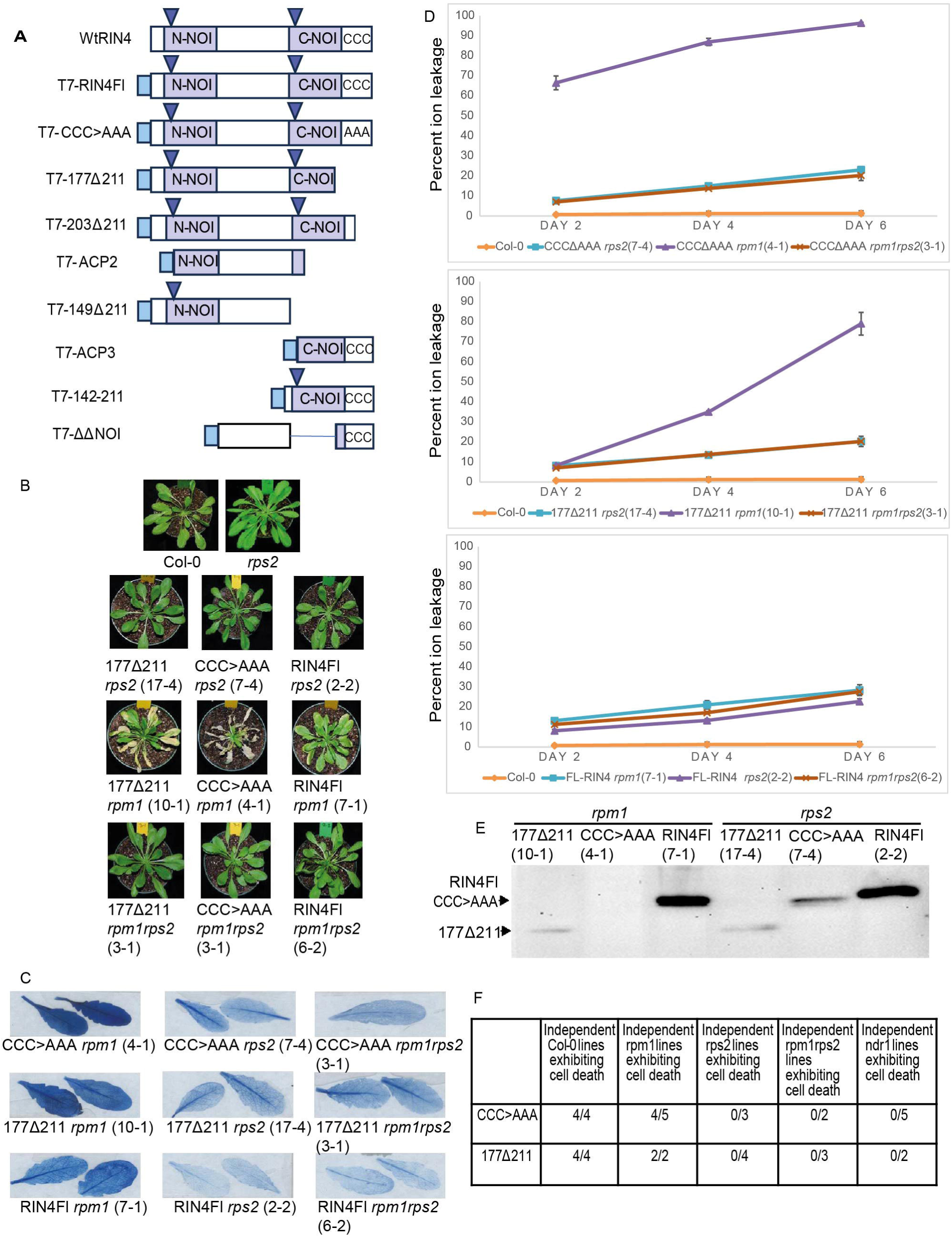
Non-membrane-tethered derivatives of RIN4 cause an RPS2-dependent cell death response in Col-0. **A)** RIN4 derivatives expressed from a Dex-inducible promoter in transgenic Arabidopsis plants. Purple rectangle, NOI-domain; Blue triangle, site of cleavage by AvrRpt2; Light blue square, T7-tag; RIN4Fl, RIN4 full length. **B)** Macroscopic symptoms were observed at 72 hours post induction (HPI) with dexamethasone (dex) of indicated RIN4 derivatives in homozygous transgenic Arabidopsis plants. **C)** Trypan blue staining of leaves from plants as in **B**. **D)** Cell death in plants as in **B** was quantified by measuring electrolyte leakage. Data was collected from 3 independent experiments with 3 technical replicates per transgenic line (n=9). Error bars represent standard error of the mean (SEM). **E)** Anti-T7 immunoblot showing protein accumulation of CCC>AAA and 177Δ211 in *rpm1* or *rps2* at 72 HPI. Arrows indicate positions of individual RIN4 derivatives. **F)** Occurrence of cell death following induction of non-membrane-tethered RIN4 derivatives in additional T2 lines of Col-0, *rpm1*, *rps2*, *rpm1rps2*, or *ndr1*.

Given its similarity to an HR, we reasoned that the cell death elicited by these non-membrane-tethered derivatives likely results from the activation of either RPM1 and/or RPS2. To test this prediction, transgenic lines expressing dex-inducible RIN4FL, 177Δ211 and CCC>AAA were established in *rpm1-3* or *rps2-101C* single mutant or *rpm1rps2* double mutant Arabidopsis plants. Similar to expression of 177Δ211 or CCC>AAA in *rpm1rps2rin4* triple mutant Arabidopsis plants ^27^, expression in either *rps2* single mutant or *rpm1rps2* double mutant plants did not elicit a cell death response **(Figures 1B C and D).** However, expression of these non-membrane tethered derivatives in *rpm1* single mutant plants elicited a cell death response **(Figures 1B C and D).** The difference in the cell death responses could not be attributed to varying accumulation levels of CCC>AAA and 177Δ211 derivatives in different transgenic lines **(Figure 1E)**. While accumulation of 177Δ211 was detectable in both *rpm1* and *rps2* plants, CCC>AAA could not be detected in *rpm1*, likely due to the stronger cell death response it causes compared to 177Δ211. We tested the response in additional, independent transgenic lines and observed that expression of CCC>AAA and177Δ211 in plants expressing RPS2 (Col-0 or *rpm1*) elicited cell death in 8 out of 9 and 6 out of 6 lines, respectively, whereas no cell death was observed in *rps2* or *rps2 rpm1* mutant lines **(Figure 1F)**. Collectively, these results indicate that activation of cell death by non-membrane-tethered RIN4 derivatives is RPS2-dependent and is not suppressed by native RIN4.

### The NOI domains mediate both suppression of RPS2 within membrane-tethered derivatives of RIN4 and activation of RPS2 within non-membrane-tethered derivatives of RIN4

Next, we sought to determine the contribution of NOI domains within membrane-tethered derivatives of RIN4, Flag-1Δ64 (lacks N-NOI), Flag-149Δ176 (lacks C-NOI) or Flag-ΔΔNOI (double deletion of 1Δ64 and 149Δ176, lacks both NOI domains), towards suppression of RPS2 **(Figure S2A).** Consistent with previous observations ^48^, Flag-1Δ64 suppressed RPS2 as effectively as Flag-RIN4FL in *N. benthamiana* **(Figure S2B)**. Despite accumulating to similar levels as FL-RIN4, Flag-149Δ176 retained modest suppressive activity whereas Flag-ΔΔNOI failed to suppress RPS2 **(Figure S2B-D)**. Thus, the NOI domains in the membrane-tethered derivatives are required for RPS2 suppression in transient assays.

To further assess the role of NOI domains within non-membrane-tethered derivatives of RIN4 in overcoming suppression of RPS2 by native RIN4, we generated plant lines that inducibly express additional derivatives of RIN4 **(Figure 1A).** Lines expressing derivatives that contain both the NOIs (CCC>AAA, 203Δ211 and 177Δ211) displayed strong cell death, while those expressing derivatives lacking the C-NOI domain displayed weaker (ACP2) or no (149Δ211) cell death **(Figures S1A-C)**. The reduced activity of ACP2 and 149Δ211 did not result from lack of expression; they accumulated to significantly higher levels than the cell-death-inducing derivatives with the C-NOI, which were barely or not detectable, likely due to the induced cell death **(Figure S1D)**. None of the plants expressing membrane-tethered derivatives of RIN4, including ACP3 (AA153-211, with a truncated C-NOI domain), 142-211 (containing an intact C-NOI domain), or ΔΔNOI (lacking both NOI domains) displayed cell death, despite readily detectable protein accumulation **(Figure S3A-D)**. However, the membrane-tethered derivatives with either a truncated (ACP3) or full (142-211) C-NOI produced chlorotic symptoms resembling those observed with RIN4FL and ACP2 **(Figure S1A and S3A)**. Collectively, data from the transient and stable transgenic assays indicate that NOI domains mediate either activation or repression of RPS2 depending on whether they are within a non-membrane-tethered or membrane-tethered derivative of RIN4, respectively.

### Activation of RPS2 by non-membrane-tethered derivatives of RIN4 requires NDR1

NDR1 is a plasma membrane-localized, glycophosphatidyl-inositol (GPI)-anchored protein required for signal transduction from several R-proteins, including RPM1, RPS2, and RPS5 ^46,52,63^. *In planta* interaction between NDR1 and RIN4 is proposed to contribute to its role in RPM1 and RPS2 signaling ^51^. The ectopic activation of RPS2 that underlies seedling lethality in homozygous *RPS2rin4* plants still occurs immediately following germination in *RPS2rin4ndr1* plants ^46^. This contrasts with activation of RPS2 by AvrRpt2, which requires NDR1.

To determine the role of NDR1 in the activation of RPS2 by non-membrane-tethered RIN4 derivatives, transgenic lines carrying inducible constructs were generated in the *ndr1-1* mutant background. Expression of CCC>AAA or 177Δ211 failed to elicit cell death in *ndr1* plants despite detectable accumulation of each derivative **(Figure 2A, 2B, 2C and 2D)**. The failure of CCC>AAA and 177Δ211 to activate RPS2 in *ndr1* mutant plants was observed in 5 and 2 independent transgenic lines, respectively **(Figure 1F)**. Thus, like the activation of RPS2 by AvrRpt2 and unlike the ectopic activation of RPS2 in the absence of RIN4, the activation of RPS2 by non-membrane-tethered derivatives of RIN4 requires *NDR1*.

**Figure 2:**
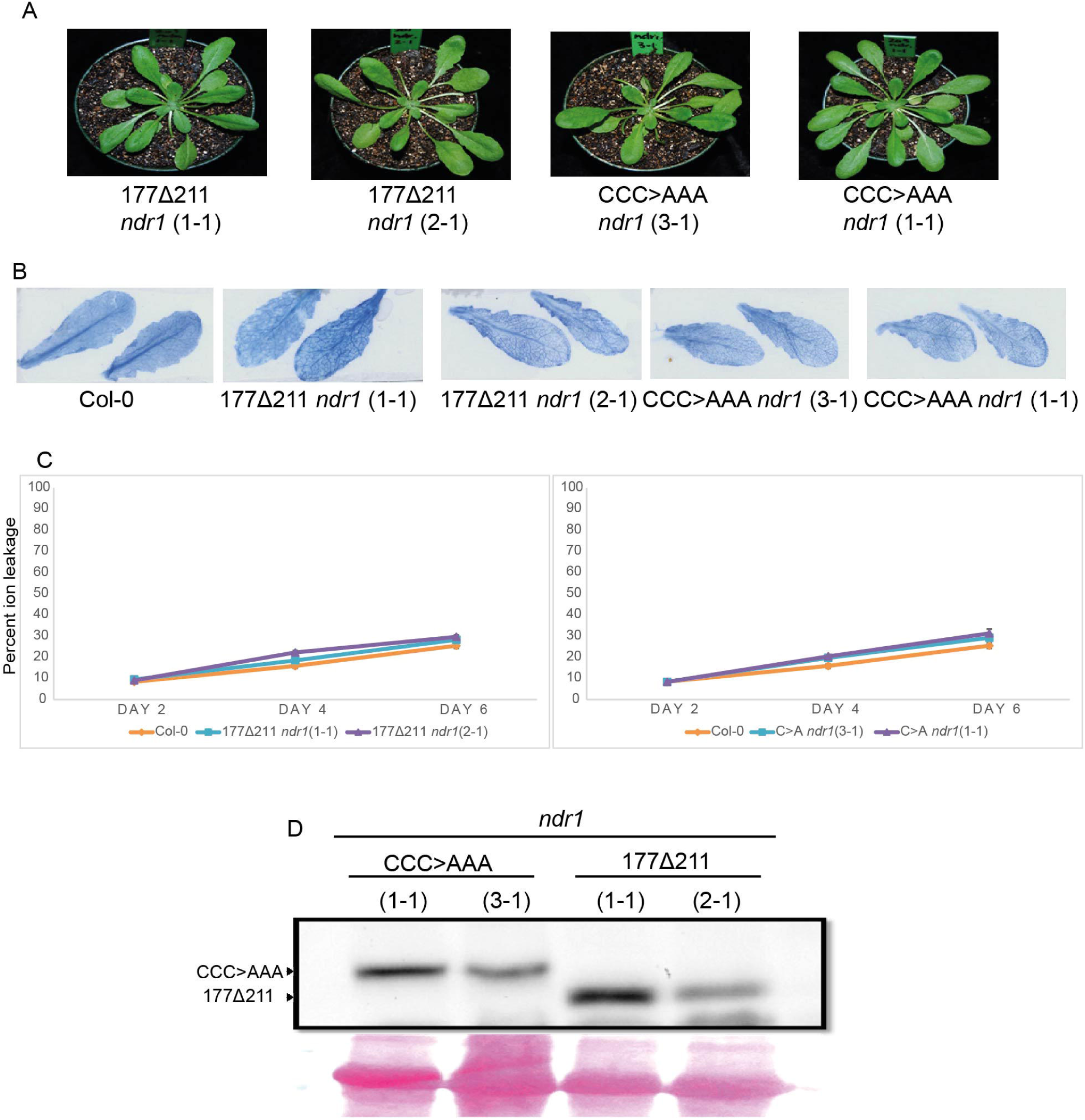
Cell death caused by the non-membrane-tethered derivatives of RIN4 is NDR1-dependent. **A)** Macroscopic symptoms were observed at 72 HPI with dex of indicated non-membrane-tethered RIN4 derivatives in homozygous transgenic *ndr1* plants. **B)** Trypan blue staining of leaves from plants as in **A**. **C)** Cell death in plants as in **A** was quantified by measuring electrolyte leakage. Data was collected from 3 independent experiments with 3 technical replicates per transgenic line (n=9). Error bars represent SEM. **D)** Anti-T7 immunoblot showing protein accumulation of CCC>AAA and 177Δ211 in *ndr1* lines at 72 HPI. The lower panel shows ponceau staining of RuBisCO as a loading control. Arrows indicate positions of individual RIN4 derivatives.

To further examine the role of AtNDR1 in ectopic and effector-induced activation of RPS2, we expressed various combinations of Flag-RIN4FL, RPS2-HA, HA-NDR1 and either AvrRpt2-HA or the proteolytically inactive mutant AvrRpt2^C122A^-HA in *N. benthamiana* leaves **(Figure S4A)**. The ectopic activity of RPS2-HA in the absence of Flag-RIN4FL was unaffected by the presence or absence of HA-NDR1 **(Figure S4B and C)**. As expected, co-expression of Flag-RIN4FL suppressed the activity of RPS2-HA and expression of AvrRpt2-HA, but not AvrRpt2^C122A^-HA, overcame the suppression and activated RPS2-HA. Interestingly, co-expression of HA-NDR1 attenuated activation of RPS2 by AvrRpt2 **(Figure S4B and C)**. Thus, as in Arabidopsis, the ectopic activity of RPS2 in *N. benthamiana* is unaffected by NDR1 ^46^. However, although constitutive overexpression of AtNDR1 did not inhibit RPS2 activation by AvrRpt2 in Arabidopsis ^63^, its transient overexpression suppresses AvrRpt2-mediated activation of RPS2 in *N. benthamiana*.

### Non-membrane-tethered derivatives of RIN4 activate RPS2 in the cytosol where they compete with intact RIN4 for association with RPS2

C-terminal acylation targets RIN4 to the plasma membrane ^23,24^. Unlike derivatives of RIN4 with an intact C-terminal acylation site, ACP2, 177Δ211 and CCC>AAA were not found in the endomembrane fraction in Arabidopsis ^27^. When expressed in *N. benthamiana*, YFP-RIN4Fl localized to the plasma membrane whereas the YFP-tagged versions of these non-membrane-tethered derivatives of RIN4 accumulated in both the cytosol and nucleus, the latter based on their co-localization with Hoechst dye **(Figure S5A-C)**.

In order to determine whether the non-membrane-tethered derivatives of RIN4 mediate RPS2 activation in the nucleus or cytosol, derivatives of CCC>AAA carrying a nuclear localization signal (NLS, YFP-NLS-CCC>AAA), a nuclear export signal (NES, YFP-NES-CCC>AAA), or a scrambled nuclear export signal (SNE, YFP-SNE-CCC>AAA) were constructed and shown to localize as expected in the nucleus, cytosol, or both, respectively **(Figure S5A and D)**. When co-expressed with RPS2 and RIN4FL, the nuclear-excluded YFP-NES-CCC>AAA derivative overcame suppression by Flag-RIN4FL and activated RPS2-induced cell death at comparable levels to ectopic activation of RPS2 in *N. benthamiana*. In contrast, nuclear-localized YFP-NLS-CCC>AAA does not activate RPS2 **(Figure S5E and F)**. This result was confirmed under conditions in which agrobacteria concentrations were adjusted to give equivalent expression of both YFP-NLS-CCC>AAA and YFP-NES-CCC>AAA **(Figure S5E-G**). Thus, cytosolic accumulation of the CCC>AAA derivative of RIN4 is required for its ability to activate RPS2.

To validate results from transient assay in *N. benthamiana*, transgenic lines of Arabidopsis (Col-0) that inducibly express CCC>AAA derivatives tagged with AcV5 and either NLS, NES, or SNE were generated **(Figure 3A)**. When individual leaves of T1 plants were treated with dexamethasone, strong cell death was observed in only 1 of 22 plants transformed with the inducible AcV5-NLS-CCC>AAA compared to 12 of 25 and 11 of 20 with the AcV5-NES-CCC>AAA or AcV5-SNE-CCC>AAA constructs, respectively **(Figure 3B)**. This observation was further supported by generating and testing three independent transgenic lines for each construct that had variable but overlapping levels of protein expression **(Figure 3C)**. Lines expressing AcV5-NES-CCC>AAA or AcV5-SNE-CCC>AAA displayed strong cell death while lines expressing AcV5-NLS-CCC>AAA did not **(Figures 3D and E)**.

**Figure 3.**
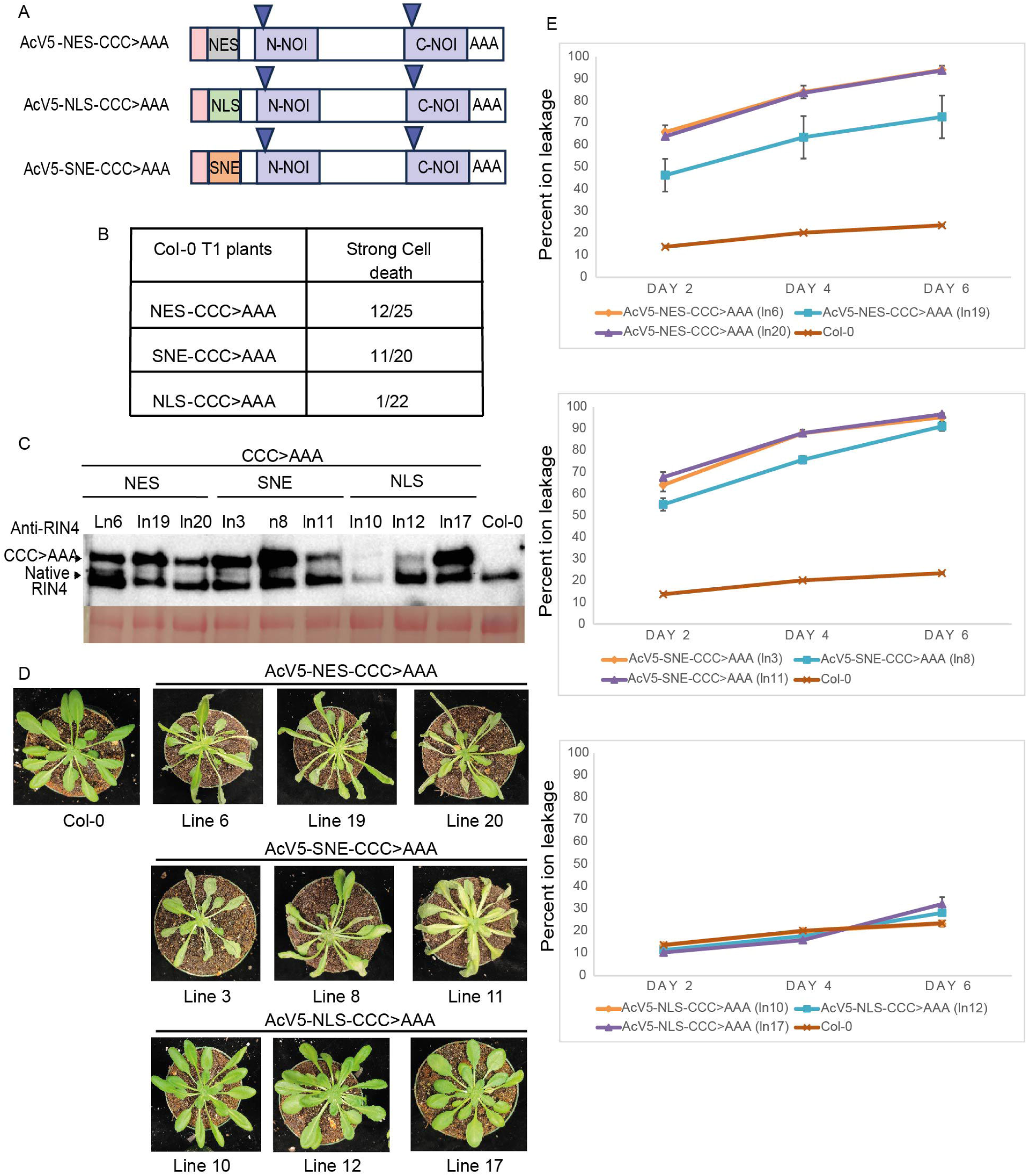
Non-membrane-tethered CCC>AAA must accumulate in the cytosol to activate RPS2 in Arabidopsis. **A)** RIN4 derivatives expressed from a Dex-inducible promoter in transgenic Arabidopsis plants. Peach rectangle, AcV5 tag; N-terminal square, NES, SNE, or NLS tags. **B)** Occurrence of symptoms at 72HPI in independent T1 lines induced to express the indicated derivatives of RIN4. **C)** Anti-RIN4 immunoblot showing protein accumulation of the RIN4 derivatives in the indicated homozygous transgenic Arabidopsis plants at 72 HPI. The lower panel shows ponceau staining of RuBisCO. Arrows indicate the positions of native RIN4 and the tagged-CCC>AAA derivatives. **D)** Macroscopic symptoms were observed at 72 HPI with dex of indicated non-membrane tethered CCC>AAA derivative in homozygous transgenic Arabidopsis. **E)** Cell death in plants as in **D** was quantified by measuring electrolyte leakage. Data was collected from 3 independent experiments with 3 technical replicates per transgenic line (n=9). Error bars represent SEM.

RIN4 and RPS2 associate *in planta* ^30,49^. To determine if RPS2 also associates with the non-membrane tethered CCC-AAA derivative, we generated 35S:RPS2-TurboID-V5 **(Figure S6A)** to enable TurboID-based proximity labelling in *N. benthamiana*. 35S:RPS2-TurboID-V5 was expressed with either Flag-RIN4FL or AcV5-NES-CCC>AAA derivative. As expected, RPS2-TurboID-V5 biotinylated Flag-RIN4FL **(Figure S6B)** ^30,49^. RPS2-TurboID-V5 also biotinylated AcV5-NES-CCC>AAA **(Figure S6B)**. The reduced labeling of AcV5-NES-CCC>AAA, relative to RIN4FL, likely results from the cell death that occurs because the AcV5-NES-CCC>AAA RIN4 derivative fails to suppress RPS2. Interestingly, labeling of Flag-RIN4FL by RPS2-TurboID-V5 was greatly diminished when AcV5-NES-CCC>AAA was co-expressed **(Figure S6C)**. Conversely, labeling of AcV5-NES-CCC>AAA was unaffected by co-expression of Flag-RIN4FL **(Figure S6D)**. These data indicate that the CCC>AAA derivative of RIN4 effectively outcompetes RIN4FL for association with RPS2. Collectively, the data indicate that non-membrane-tethered derivatives of RIN4 activate RPS2 at the plasma membrane, possibly by displacing association of RPS2 with inhibitory, membrane-tethered RIN4.

### ACP2 associates with RPS2 to overcome ACP3-mediated suppression and activate RPS2

Collectively, our data indicate that NOI domains contribute to inhibition or activation of RPS2 depending on whether they are in a membrane-tethered or cytosolic form. Furthermore, RPS2 activation by cytosolic NOI domains is dependent on NDR1. These findings led us to speculate that, during the NDR1-dependent activation of RPS2 by AvrRpt2, suppression of RPS2 by ACP3 is overcome by ACP2. To test this hypothesis, we co-expressed Flag-ACP2 and/or Flag-ACP3 along with RPS2-HA in *N. benthamiana* (**Figure 4A)**. Although Flag-ACP3 was expressed to lower levels than RIN4FL, both were equally competent to suppress RPS2-HA (**Figure 4B, C and D**). As expected, Flag-ACP2 failed to suppress RPS2-HA. Consistent with our hypothesis, co-expression of Flag-ACP2 overcame suppression by Flag-ACP3 and activated RSP2-HA. Furthermore, as was observed for RPS2 activation by AvrRpt2, overexpression of NDR1-HA suppressed the ability of Flag-ACP2 to overcome suppression by Flag-ACP3 and activate RPS2-HA **(Figure S4D and E)**.

**Figure 4.**
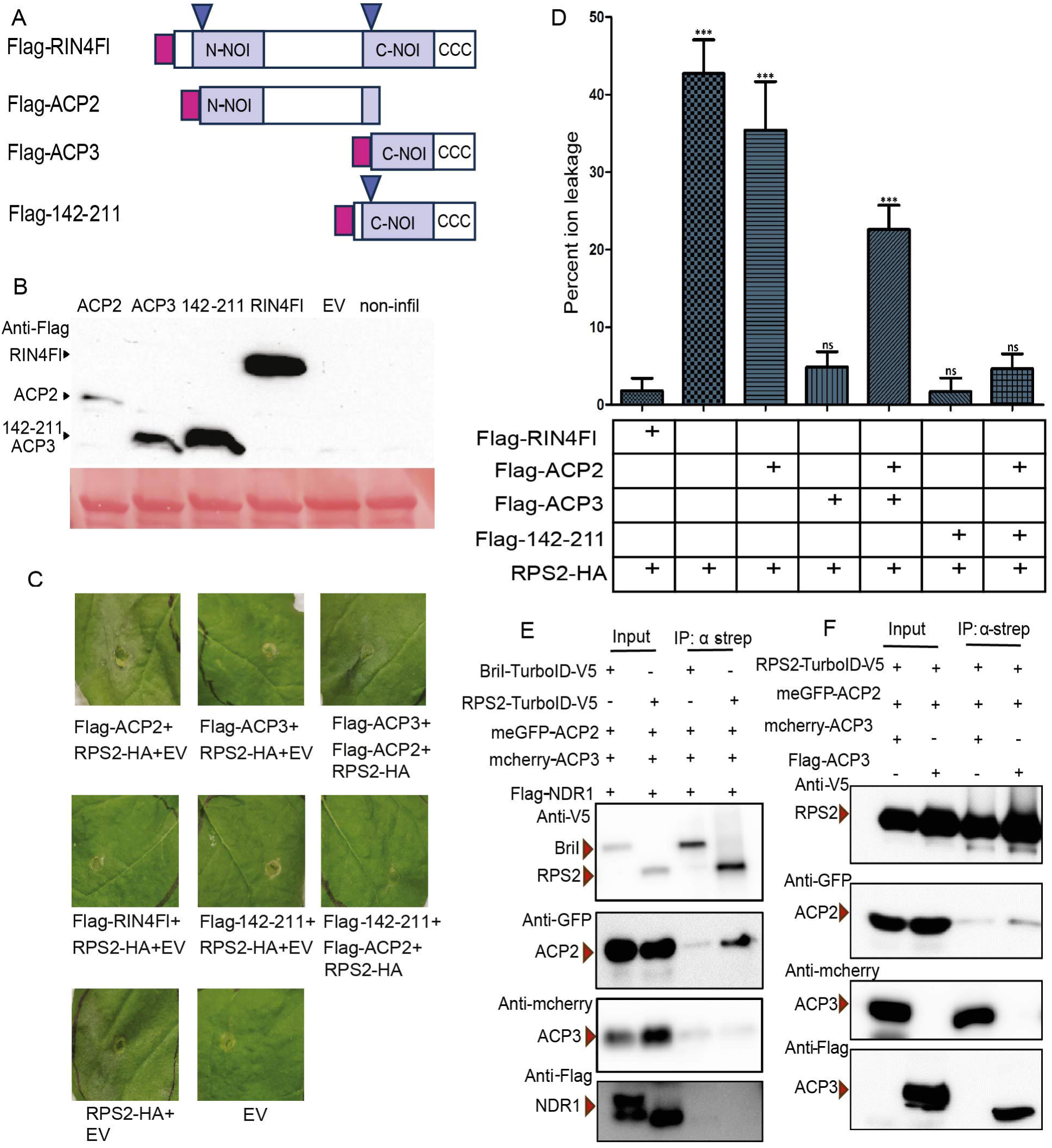
**ACP2 activates RPS2 suppressed by ACP3 in *N. benthamiana***. **A)** RIN4 derivatives expressed from 35S promoter in transient experiments in *N. benthamiana*. Pink rectangle, Flag-tag. **B)** Anti-Flag immunoblot showing protein accumulation of the indicated RIN4 derivatives in *N. benthamiana* at 72 HPI. The lower panel shows ponceau staining of RuBisCO. Arrows indicate position of individual RIN4 derivatives. **C)** Macroscopic symptoms at 48 hours after infiltration (HAI) for agro-transient expression of RIN4 derivatives (each at OD_600_ = 0.6) and RPS2-HA (at OD_600_ = 0.04) of *N. benthamiana* leaves. **D)** Cell death in plants as in **C** was quantified by measuring electrolyte leakage at 96 HAI. Data was collected from 3 independent experiments with 3 technical replicates per treatment (n=9). Error bars represent SEM. Student’s t-test, at 95% confidence limits, was used for comparison with combination of RPS2-HA and Flag-RIN4Fl (ns, not significant; ***P<0.0001). **E)** *N. benthamiana* leaves were infiltrated with agrobacterium containing the indicated constructs (each at OD_600_ = 0.5). After 24hrs the leaves were infiltrated with biotin and 4hrs later tissue was collected. Input and anti-strep IP samples were immunoblotted as indicated with red arrows indicating the position of the proteins. **F)** *N. benthamiana* leaves were co-infiltrated with agrobacterium containing the indicated RIN4 derivatives and either RPS2-TurboID-AcV5 (each at OD_600_ = 0.5) and processed as in **E**.

Since ACP2 accumulates in both the cytosol and nucleus ^27^, we sought to confirm that cytosolic ACP2 activates RPS2. Derivatives of ACP2 with YFP and either NLS, NES, or SNE at their N-termini localized as expected when expressed in *N. benthamiana* (**Figure 5A and 5B**). When these derivatives were expressed in *N. benthamiana* along with Flag-ACP3 and RPS2-HA, YFP-SNE-ACP2 or YFP-NES-ACP2, but not by YFP-NLS-ACP2, activated RPS2-HA **(Figures 5C and D)**. Expression levels of the proteins do not account for their (in)ability to activate ACP3-suppressed RPS2 **(Figure 5D)**. Thus, as for non-membrane-tethered derivatives of RIN4, ACP2 must be present in the cytosol to activate RPS2.

**Figure 5.**
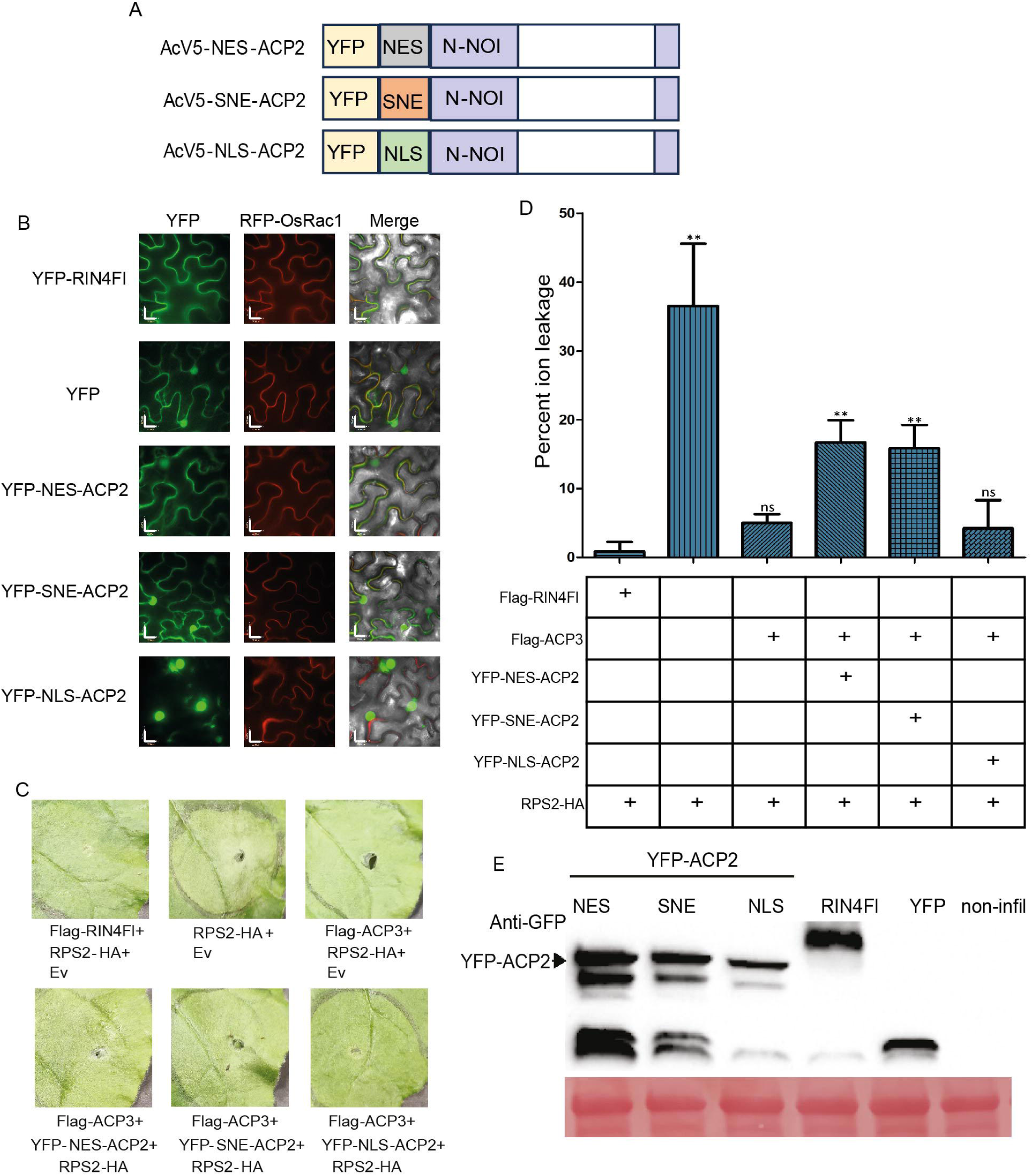
ACP2 must accumulate in the cytosol to activate RPS2 in *N. benthamiana*. **A)** ACP2 derivatives expressed from 35S promoter in transient experiments in *N. benthamiana*. AcV5 tags are followed by rectangles indicating, NLS, NES, or SNE tags. **B)** The indicated derivatives of RIN4 were co-expressed with the plasma membrane marker RFP-OsRac1 in *N. benthamiana*. Shown are YFP, RFP and merged confocal microscope images at 72 HAI. Scale bars are 25 µM. **C)** Macroscopic symptoms at 48 HAI for agro-transient expression of RIN4 derivatives (OD_600_ = 0.6) and RPS2-HA (OD_600_ = 0.04) of *N. benthamiana* leaves. **D)** Cell death in plants as in **C** was quantified by measuring electrolyte leakage at 96 HAI. Data was gathered from 4 independent experiments with 3 technical replicates per treatment (n=12). Error bars represent SEM. Student’s t-test, at 95% confidence limits, was used for comparison with combination of RPS2-HA and Flag-RIN4Fl (ns, not significant; **P<0.01). **E)** Anti-GFP immunoblot showing accumulation of the indicated proteins in *N. benthamiana* at 72 HAI. Arrows indicate position of the RIN4 derivatives.

ACP2 associates with RPS2 dependent on access enabled by cleavage within the C-NOI. Similar to Flag-ACP3, a derivative of RIN4 with an intact C-NOI domain, Flag-142-211 also suppressed RPS2 in *N. benthamiana* (**Figure 4A)**. However, Flag-ACP2 failed to activate RPS2-HA suppressed by Flag-142-211 (**Figure 4C and D**). This difference likely results from steric interference rather than a difference between an intact and cleaved C-NOI since Flag-ACP2 also failed to activate RPS2-HA suppressed by ACP3 with a bulky C-terminal adduct, YFP-ACP3 **(Figure S7A and B)**. Also, similar to its weak activation of RPS2 suppressed by native RIN4 in Arabidopsis transgenic lines **(Figures S2)**, Flag-ACP2 failed to activate RPS2-HA that was suppressed by Flag-RIN4FL in *N. benthamiana* **(Figure S7C and D).** We additionally determined whether RPS2-TurboID-V5 is able to biotinylate ACP2 in the presence of RIN4Fl. To facilitate protein accumulation, EST:RPS2-TurboID-V5 construct was used for this assay. Co-expression of Flag-RIN4Fl and meGFP-ACP2 with EST:RPS2-TurboID-V5, resulted in biotinylated of RIN4Fl specifically by RPS2-TurboID-V5 (**Figure S7E**). This result indicated that Flag-RIN4Fl out-competes meGFP-ACP2 for binding to RPS2.

These data support the hypothesis that ACP2 activates RPS2 dependent on access to an ACP3-RPS2 complex created upon cleavage of RIN4 within its C-terminal NOI.

We further tested this hypothesis by using TurboID to examine the proximity of ACP2 to RPS2 under various conditions. To facilitate protein accumulation before the onset of RPS2 induced HR, EST:RPS2-TurboID-V5 construct was used for this assay. When co-expressed with mcherry-ACP3, Flag-NDR1, and either EST:Bri1-TurboID-V5 or EST:RPS2-TurboID-V5, meGFP-ACP2 was biotinylated specifically by RPS2-TurboID-V5 (**Figure 4E**). Surprisingly, labeling of neither Flag-NDR1 nor mcherry-ACP3 was detected. Thus, even when ACP2 is unable to activate RPS2 because it is suppressed by ACP3 with a bulky C-terminal adduct, it is still in close proximity to the ACP3-RPS2 complex. The biotinylation of meGFP-ACP2 by RPS2-TurboID-V5 was stronger in the presence of Flag-ACP3 than mCherry-ACP3, despite the activation of cell death in the latter sample (**Figure 4F**). The presence or absence of Flag-NDR1 in these experiments had distinct effects on the labeling of ACP2 and ACP3. Flag-NDR1 enhanced labeling of meGFP-ACP2 (compare strong signal in 4E relative to that in 4F with mcherry-ACP3). Conversely, it prevented labeling of Flag-ACP3 (compare strong signal in 4F to the lack of specific labeling in 4E). Collectively, these results indicate that ACP2 is recruited to the RPS2 complex by NDR1 and most efficiently accesses the complex when the N-terminus of ACP3 is exposed.

### Activation of RPS2 requires compatibility between ACP2 and ACP3

We previously demonstrated that C-terminal fragments equivalent to ACP3 (YFP-RIN4^CLV3^, referred to as YFP-ACP3 in the current study) from RIN4 homologs of soybean, apple, peach, and potato, but not rice, are capable of suppressing RPS2-HA when transiently expressed in *N. benthamiana* ^64^. To examine the ability of ACP2-equivalent fragments to activate RPS2 that was suppressed by RIN4 homologs from other plants, we generated Flag-tagged versions of ACP3 without the bulky, N-terminal YFP-tag. As for the YFP-tagged versions, Flag-ACP3 derivatives from soybean, peach, or potato, but not rice, suppressed RPS2 **(Figure 6A, B, C and S8)**. As expected, RPS2-HA suppressed by Flag-AtACP3, but not YFP-AtACP3, was activated by Flag-AtACP2 as well as a related derivative, YFP-AtACP2, that lacks ACP1 and the N-terminal RIN4 cleavage site **(Figure 6D).** Surprisingly, YFP-AtACP2 failed to activate RPS2-HA that was suppressed by the Flag-ACP3 derivatives from soybean, peach, or potato. However, RPS2-HA suppressed by these Flag-ACP3 derivatives was activated by the YFP-ACP2 derivatives from the corresponding plant species. The necessary compatibility between ACP2 and ACP3 indicates that, following their generation upon cleavage by AvrRpt2, these fragments may directly interact during activation of RPS2.

**Figure 6.**
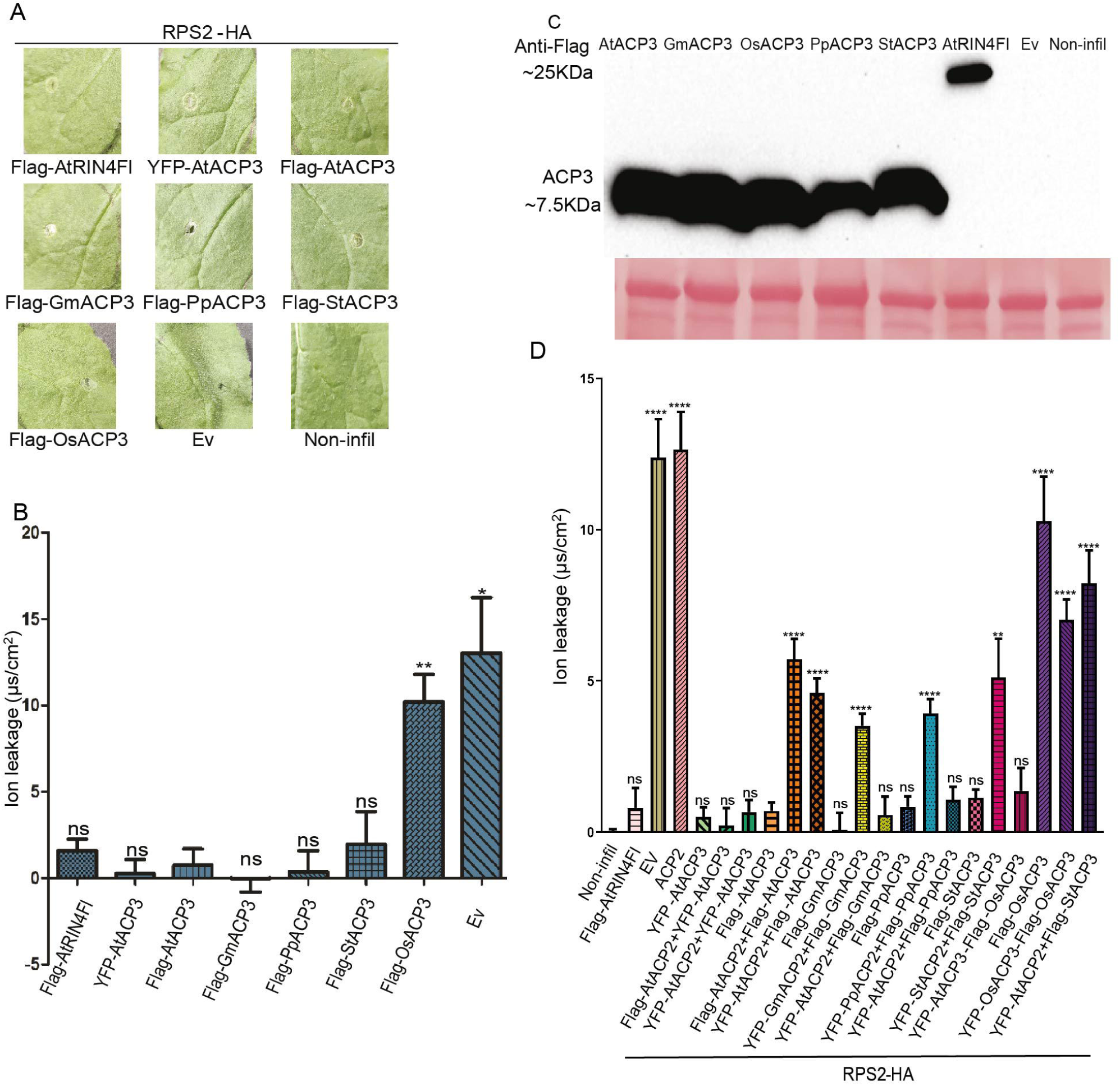
Activation of RPS2 suppressed by ACP3 requires compatibility between ACP2 and ACP3. **A)** Macroscopic symptoms at 48 HAI for agro-transient expression of RIN4 derivatives (OD_600_ = 1.0) and RPS2-HA (OD_600_ = 0.05) of *N. benthamiana* leaves. **B)** Cell death in plants as in **A** was quantified by measuring electrolyte leakage at 72 HAI. Data was collected from 3 independent experiments with 3 technical replicates per treatment (n=9). Error bars represent SEM. Student’s t-test, at 95% confidence limits, was used for comparison with Flag-AtACP3 (ns, not significant; *P<0.05, **P<0.01). **C)** Anti-Flag immunoblot shows accumulation of Flag-RIN4 derivatives expressed in *N. benthamiana* at 72 HAI. The lower panel shows ponceau staining of RuBisCO. **D)** Electrolyte leakage at 96 HAI for agro-transient expression of RIN4 derivatives (each OD_600_ 0.6) and RPS2-HA (OD_600_ 0.04) in *N. benthamiana* leaves. Data was collected from 5 independent experiments with 3 technical replicates per treatment (n=15). Error bars represent SEM. Student’s t-test, at 95% confidence limits, was used for comparison with Flag-AtACP3 (ns, not significant; **P<0.0042; ****P<0.0001).

### Spatial proximity of RPS2 and ADR1-L1 in *N. benthamiana*

Previous studies have reported that in *A*. *thaliana*, ADR1 regulates and induces ETI responses downstream of many sensor NLRs ^60–62^. Additionally, it was reported that in *adr1* triple mutant plants, activation of RPS2 by AvrRpt2 is significantly attenuated ^60^. *ADR1* genes thus function to support ETI mediated by activated RPS2. Sensor NLR specificity for a particular helper NLR may or may not require physical association of the sensor and helper NLRs. Based on the involvement of the helper ADR1 in mediating RPS2-triggered ETI responses ^60^ we speculated that activated RPS2 might interact with ADR1. To test RPS2-ADR1 interactions, co-immunoprecipitation experiments were performed following co-expression of EST:RPS2-TurboID-V5 with 35S:ADR1-L1-Flag. Unlike the EST:BRI1-TurboID-V5 control, EST:RPS2-TurboID-V5 weakly co-immunoprecipitated with 35S;ADR1-L1-Flag **(Figure S9A)**. To further test this interaction, we used Turbo-ID proximity labelling. Surprisingly, 35S:ADR1-L1-Flag was only weakly biotinylated by EST:RPS2-TurboID-V5 and was more strongly biotinylated by EST:Bri1-TurboID-V5 **(Figure S9B)**. We conclude that since RPS2-TurboID-V5 gives no more biotinylation of ADR1-L1 than does BRI1-Turbo-ID, the contribution of ADR1 to RPS2 signaling is unlikely to be via direct interaction.

## Discussion

NLRs activate defense upon effector recognition by diverse mechanisms. Within a phylogenetic tree of CNLs, RPS2 (and the related RPS5) are part of a distinct G10 clade ^18^. The mechanisms by which effector-dependent perturbation of RIN4 activates RPS2, and other CNLs that “guard” RIN4, has remained obscure.

Here we demonstrate that non-membrane-tethered derivatives of RIN4 activate RPS2 via a mechanism that is distinct from its ectopic activation in the *absence* of RIN4 **(Figure 7A and B)**. Furthermore, translating these findings to the activation of RPS2 by AvrRpt2, we demonstrate coordinated roles for ACP2 and ACP3, the AvrRpt2-cleavage fragments of RIN4. While ACP3, along with NDR1, maintains RPS2 within a pre-activation complex **(Figure 7C)**, ACP2 interacts with the pre-activation complex to activate RPS2 **(Figure 7D)**. Triggering of RPS2 in the *presence* of ACP2 is reminiscent of triggering of the apple NLR, Mr5, in the *presence* of ACP3. Collectively, these findings support the model in which cleavage of RIN4 by AvrRpt2 generates both ACP3 with a truncated C-NOI that primes RPS2 and ACP2 that triggers RPS2 activation by interacting with the NDR1-ACP3-RPS2 pre-activation complex.

**Figure 7.**
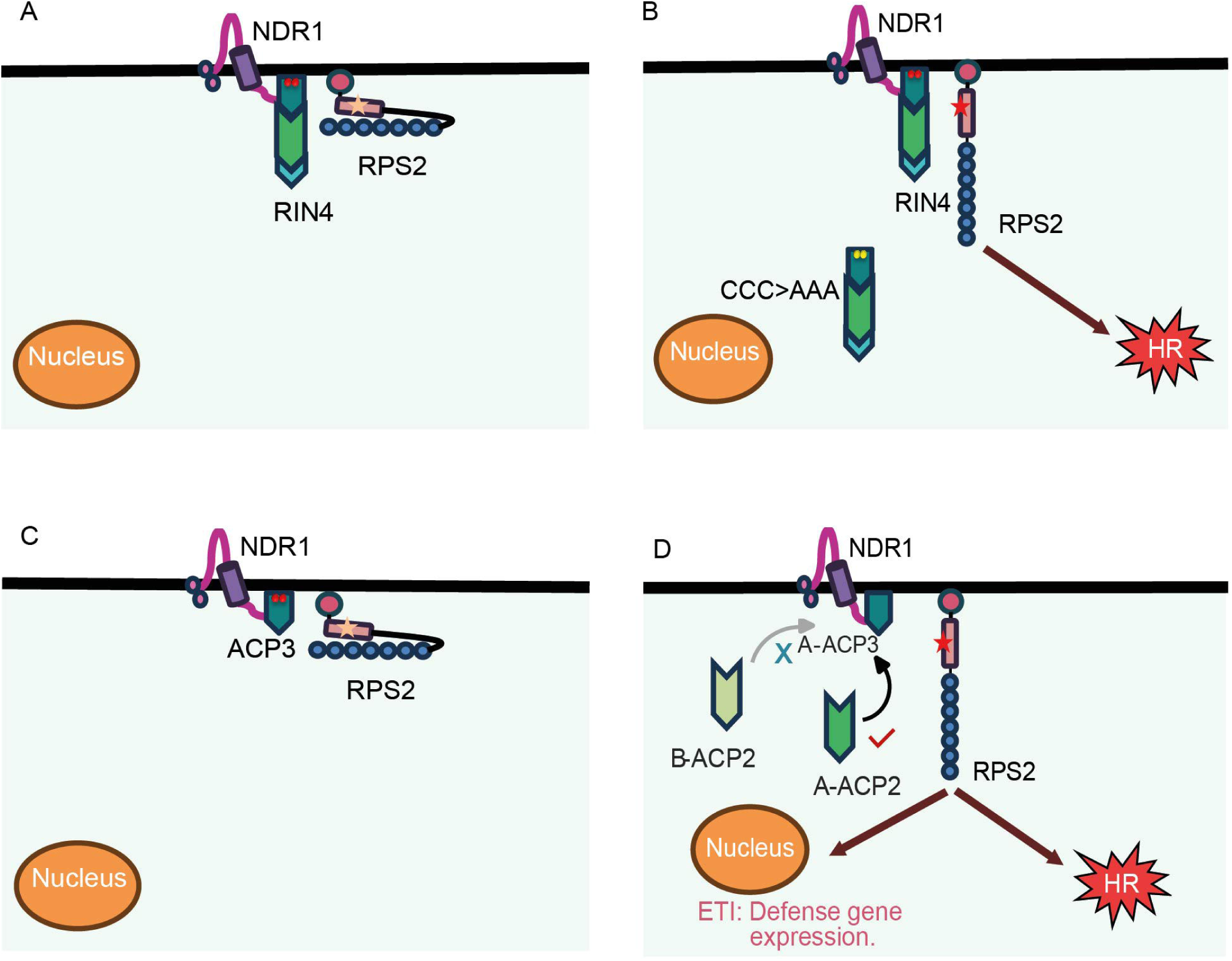
Model of RPS2 resting, pre-activation and activation complexes. **A)** In an unstimulated cell, the two membrane-tethered NOI domains of RIN4 maintain RPS2 in a resting complex that also includes NDR1. **B)** A non-membrane-tethered derivative of RIN4 with two NOI domains, *e.g.*, CCC>AAA, activates RPS2 dependent on NDR1. **(C-D)** The AvrRpt2 cleavage products of RIN4, ACP2 and ACP3, play contrasting roles in RPS2 regulation. **C)** ACP3 maintains RPS2 in a pre-activation complex with NDR1. **D)** ACP2 interacts with the complex to trigger activation of RPS2. Activation requires both NDR1 and compatibility between ACP2 and ACP3. For example, RPS2 suppressed by ACP3 from plant species “A” is activated by ACP2 from plant species “A” but not by ACP2 from plant species “B”.

Our model is supported by examining regulation of RPS2 by RIN4 from other angiosperms. RPS2 suppression by ACP3 derivatives from soybean, peach, and potato is relieved only by co-expression of ACP2-like fragments from the same plant species, but not by the comparable fragment from Arabidopsis. Thus, the C- and N-NOI domains of RIN4 appear to have co-evolved to maintain functional compatibility, perhaps via their direct interaction within the RPS2 activation complex. The idea that the fragments of RIN4 make distinct, cooperative contributions during RPS2 activation is supported by the greater level of similarity within than between C-NOIs and N-NOIs of diverse plant species^27^.

Our model is further supported by the observation that activation of RPS2 by non-membrane-tethered derivatives of RIN4 requires *NDR1*, which aligns with the requirement of *NDR1* for RPS2 activation by AvrRpt2 and is distinct from the NDR1-independent ectopic activation of RPS2 in *rin4* plants. Surprisingly, AtNDR1 expression in *N. benthamiana* inhibited AvrRpt2-mediated activation of RPS2. A recent study reported that overexpression of NDR1 in Arabidopsis results in interaction of NDR1 with RIN4 which protects it from being cleaved by AvrRpt2 ^65^. We speculate that overexpression of NDR1 in *N. benthamiana* might suppress RPS2 activation by stabilizing the putative pre-activation complex consisting of RPS2, NDR1, and intact RIN4. Moreover, AtNDR1 expression also suppressed ACP2-mediated activation of ACP3-inhibbited RPS2, suggesting AtNDR1 may have a suppressive function distinct from or additional to preventing cleavage of RIN4.

Inappropriate activation of NLRs in the absence of elicitation is detrimental to plant health. Thus, NLRs must be held in a tightly inactive pre-activation state. Our findings indicate that each both NOI domains of membrane-tethered RIN4 contribute to keeping RPS2 inactive, with C-NOI playing a dominant role. While ACP2 can fully activate RPS2 suppressed by ACP3, that has a truncated C-NOI domain, it fails to activate RPS2 suppressed by 142-211, with an intact C-NOI, or RIN4FL, with both NOI domains. The contribution of both NOI domains of membrane-tethered RIN4 in holding RPS2 in an inactive state is mirrored by the contribution of both NOI domains of non-membrane-tethered derivatives in overcoming that suppression. The ability of non-membrane-tethered derivatives of RIN4 with both NOI domains (CCC>AAA or 177Δ211) to overcome the strong suppression of RPS2 by RIN4 is contrasted by the inability of RIN4 derivatives with only a single NOI domain (149Δ211 and ACP2) to do so. Thus, the predominant state of RIN4 in an unchallenged plant, *i.e.* membrane-tethered and containing two NOI domains, is likely a key contributor to the minimal activity of RPS2 in the absence of AvrRpt2.

In addition to being tightly suppressed in the absence of activation, NLRs must be specifically and rapidly activated to effectively confer resistance. This can be achieved through formation of pre-assembled complexes responsive to perturbation by an effector. At the plasma membrane, RIN4 associates with both RPS2 and NDR1. While ACP3 can suppress RPS2 in transient assays, it is also part of a pre-activation complex consisting of RPS2, NDR1, and ACP3 ^25,51^. Furthermore, the partially truncated C-NOI within ACP3 is essential for triggering of the pre-activation complex by ACP2; ACP2 was unable to activate RPS2 suppressed by a derivative of RIN4 with an intact C-NOI (142-211) or a derivatives of ACP3 with a bulky C-terminal adduct. Although ACP2 and other non-membrane-tethered fragments of RIN4 accumulate both in the cytosol and the nucleus, molecules in the nucleus are unable to activate RPS2 and the required compatibility between ACP2 and ACP3 indicates their likely interaction at the plasma membrane. Thus, when RIN4 is cleaved by AvrRpt2, components previously known (NDR1) and shown here (ACP2 and ACP3) to participate in RPS2 activation are already or immediately present.

ACP3 suppression of RPS2 contrasts with the regulation of the NLR MR5 in apple, which also responds to AvrRpt2-induced cleave of MdRIN4 but is not ectopically active ^42^. Instead, AvrRpt2-induced MdACP3 fragment is both necessary and sufficient for MR5 activation, highlighting distinct mechanisms shaped by independent NLR evolution ^42^. Notably, each model posits that the virulence activity of AvrRpt2, manifested through proteolysis of RIN4, produces the trigger for NLR activation. Recently another independently evolved NLR, Ptr1 from tomato, was demonstrated to respond to AvrRpt2-induced cleavage of SlRIN4 ^43^. It will be interesting to determine the role of the AvrRpt2-induced fragments of SlRIN4 in the activation of Ptr1.

AvrRpt2 promotes *P. syringae* growth in plants lacking RIN4, indicating that targets additional to RIN4 can contribute to its virulence activity ^66^. Candidates for these virulence targets include a family of Arabidopsis proteins containing a single NOI domain, which are known to be proteolytically targeted by AvrRpt2 ^38,67^. Given that these “single-NOI” proteins also share a similar C-terminal acylation motif with RIN4, their cleaved products may share a similar activity with ACP3, which we have shown is a potent suppressor of Arabidopsis defense ^27^. Non-membrane-tethered ACP2 is also a potent suppressor of Arabidopsis defense but, unlike ACP3, it is also able to suppress flg22-induced callose deposition ^27^. Whether the nuclear localization of ACP2 contributes to its ability to suppress plant immunity remains unclear. In either case, AvrRpt2 may make a quantitatively stronger and distinct contributions to bacterial virulence by targeting RIN4, relative to single-NOI proteins. Thus, RPS2 may have evolved to “guard” RIN4, rather than a single-NOI protein, because its mechanism for perception of AvrRpt2 depends on production of the locally-concentrated, but non-membrane-tethered, NOI cleavage product, ACP2. This model is consistent with the prediction of the guard hypothesis that it is the virulence-promoting perturbation produced by AvrRpt2 that triggers RPS2.

Lastly, we investigated the putative interaction of RPS2 and ADR1 *in planta*. The two proteins were weakly associated in a co-immunoprecipitation assay, but testing of their spatial proximity revealed surprisingly low biotinylation of ADR1-L1 by RPS2-TurboID-V5. Based on our results and previous reports we speculate that ADR1-mediated potentiation of the defense response induced by activated RPS2 does not involve direct RPS2/ADR1 interactions. ADR1 supports the early steps of transcriptional reprogramming and the timely activation of HR during RPS2-mediated ETI ^60,62^. Additionally, ADR1 might contribute quantitatively to RPS2 mediated immunity by regulating defense gene expression **(Figure S10)**. It is puzzling that mutations in EDS1 have a negligible effect on RPS2 function ^68^ even though RPS2 requires ADR1 family proteins for full function, since their function usually involves formation of a heterotrimer with PAD4 ^69,70^. This raises the possibility that ADR1 might also make EDS1-independent contributions to plant immunity. Overall, our work provides valuable insights into the molecular mechanism of the AvrRpt2-RIN4-RPS2 defense activation module. The molecular details of how activated RPS2 mediates downstream signaling and interacts with ADR1 remain unclear and require further investigation.

Data availability

All data associated with the study are in the paper or supplemental information and can also be requested from the lead contact.

## Methods

### Experimental models and details

#### *Arabidopsis thaliana* and *Nicotiana benthamiana*

*Arabidopsis thaliana* (Col-0, transgenic and mutants) and *Nicotiana benthamiana* plants were grown in potting soil under a cycle of 8-h-light at 23°C and 16-h-dark at 16°C.

##### Escherichia coli

*E. coli* (Dh5 or Top10) carrying the respective constructs were grown at 37°C in Lb medium containing the appropriate antibiotic.

##### Agrobacterium tumefaciens

*Agrobacteria tumefaciens* (strain GV-3101) carrying expression constructs or empty vectors were grown overnight at 28°C in LB medium containing appropriate antibiotics (Rifampicin, Gentamicin and Kanamycin, Spectinomycin or Tetracycline).

### Method details

#### Constructs

All constructs for RIN4 derivatives (Figures 1A) were derived from pMAC100c vector containing full-length RIN4 coding sequence ^27^. These derivatives include the CCC>AAA (mutation of acylation site cysteines to alanines), 177Δ211 (deletion of 35 C-terminal residues of RIN4), 11-211 (Deletion of ACP1), ACP2 (11-152), ACP3 (153-211), 142-211 (similar to ACP3, but contains the full C-NOI), ΔΔNOI (deletion of 1Δ64 and 149Δ176), 203Δ211 (deletion of 8 C-terminal residues of RIN4), and 149Δ211 (deletion of 62 C-terminal residues of RIN4). The RIN4 derivatives were fused with an N-terminal epitope tag [T7 (MASMTGGQQMG), ACV5 (SWKDASGWS), or Flag (DYKDDDDK)] or with yellow florescent protein (YFP)]. For localization studies, Nuclear Localization Signal (NLS, PKKKRKVED) ^71^, Nuclear Export Signal (NES, NELALKLAGLDINKT) ^72^, or Shuffled Nuclear Export signal (SNE, NELALKAAGADINKT) tags were added between the N-terminal florescent/epitope tags and the C-terminal sequences encoding RIN4 or its derivatives. For transient studies in *N. benthamiana*, these chimeric fragments were cloned into pENTR-D-TOPO and subsequently moved into the gateway binary vectors pB2GW7or pGWB12 (containing a 35S promoter) or pEarlyGate 104 (35S:N-YFP).

Both 35S promoter driven (35S:) and estradiol inducible (Est:) RPS2-Turbo-ID-V5 and BRI1-Turbo-ID constructs were generated through golden gate cloning ^73^. YFP-RIN4Fl, YFP-AcV5-INT (internal) and YFP-ACP3 expressed under the control of a 35S promoter (pEarley gate 104 vector backbone) have been described previously ^64^. RPS2-HA, expressed under the control of a strong promoter (in pOCS), and HA-NDR1 have been described previously ^48,51^. 35S:AvrRpt2-HA and 35S:AvrRpt2^C122A^-HA have been described previously ^42^. 35S:RFP-OsRac1 (pGDR vector backbone), used as a plasma membrane marker, has been described previously ^74^.

### Plant Transformations

Transgenic Arabidopsis lines expressing dexamethasone (dex)-inducible tagged derivatives of RIN4 were generated as previously described ^27^ in Col-0, *ndr1-1, rpm1-3, rps2-101c* or *rpm1rps2* backgrounds. Transgene expression was induced by painting single leaves of T1 plants or spraying T2 and T3 plants with a solution containing 20 μM dex (Sigma-Aldrich, Saint Louis, MO) and 0.05% Silwet L-77 (Momentive, Waterford, NY). T1 families showing a segregation ratio of 3:1 for resistance:susceptibility to hygromycin were self-fertilized and propagated to homozygosity.

### Transient Expression Assays

*Agrobacteria tumefaciens* (strain GV-3101) carrying expression constructs or empty vectors were grown overnight at 28°C in LB medium containing appropriate antibiotics (Rifampicin, Gentamicin and Kanamycin, Spectinomycin or Tetracycline). Cultures were centrifuged at 4,000 rpm for 20 minutes and the cell pellets were subsequently re-suspended in infiltration buffer (10 mM MgCl_2_, 10 mM MES, and 200μM Acetosyringone adjusted to pH 5.6 with KOH). For all infiltration experiments the OD_600_ of cultures was adjusted with infiltration buffer and when necessary, the final OD_600_ was held constant by supplementing with bacteria carrying an empty vector. Final cultures were infiltrated into *N. benthamiana* leaves as described previously ^48,75^.

### Immunoblot Analysis

Immunoblot analysis was performed as described previously ^27^. Briefly, plant material was homogenized in a buffer containing 20 mM Tris-HCl [pH 7.5], 150 mM NaCl, 1 mM EDTA, 1 % Triton X-100, 0.1 % SDS, 5 mM DTT and 1X plant protease inhibitor cocktail and centrifuged at 20,000 × g for 10 minutes at 4°C. The concentration of protein in the supernatant was determined by the Bio-Rad protein assay reagent (Bio-Rad, Hercules, CA) according to manufacturer’s instructions. Anti-RIN4 sera (Mackey et al., 2002), anti-T7 monoclonal antibody (Novagen, Madison, Wisconsin), anti-AcV5 antibody (eBioscience, San Diego, CA), anti-Flag antibody (Sigma-Aldrich, Taufkirchen, Germany) and anti-YFP antibody (Abcam, Cambridge, UK) were used at dilutions of 1:5000, 1:10,000, 1:5000, 1:5000 or 1:5000, respectively. Chemiluminescent detection and band quantification were done using the ChemiDoc XRS system (Bio-Rad, Hercules, CA) and ImageJ (imagej.nih.gov/ij/).

### TurboID-based immunoprecipitation (IP)

For TurboID-IP assays with either 35S:RPS2-TurboID-V5 or 35S:Bri1-TurboID-V5 and additional constructs were transiently expressed in four-week-old *N. benthamiana* leaves. After 24 hours (hrs), the infiltrated leaves were treated with 50 µM biotin for 2 hrs. The leaves were then ground to a fine powder in liquid nitrogen and homogenized in extraction buffer (Glycerol 10%, 150 mM Tris-HCl [pH7.5], 1mM EDTA and 150mM NaCl, 10 mM DTT, 0.4% Nonidet-40 (Igepal), Anti-protease cocktail, 2% PVPP and 0.5% sodium deoxycholate (w/v)). The supernatant from homogenized samples was centrifuged three times at 5000 g at 4°C for 10 mins. To remove cell wall debris the supernatant was passed through miracloth into Zeba™ Spin Desalting Columns (Thermo Fisher Scientific, Catalog number 89893) and subsequently centrifuged for 1 min at 4°C to remove excess biotin. 120 µl of flowthrough was collected as input, while the rest was incubated at 4°C with 30 µl streptavidin-coated beads (Pierce™ High Capacity Streptavidin Agarose, Thermo Fisher, Catalogue number 20361). After 2 hrs, the beads were spun down at 5000 g at 4°C for 3 mins and washed 4 times with washing buffer (Glycerol 10%, 150 mM Tris-HCl [pH7.5], 1mM EDTA and 150mM NaCl, 10 mM DTT, 0.4% Nonidet-40 (Igepal), Anti-protease cocktail and 0.5% sodium deoxycholate (w/v)). Finally, 3X SDS loading buffer (3.3% SDS, 94 mM Tris-HCl pH 6.8, 30% glycerol and 0.05% (vol/vol) bromophenol blue) was added to beads and input samples followed by boiling at 95°C for 7 min to denature proteins. Denatured proteins were analysed by immunoblots.

TurboID-IP assays with Est:RPS2-TurboID-V5 or Est:Bri1-TurboID-V5 were done as described above. However, the leaves were treated with 50 µM biotin and 50 nM estradiol 48 hrs post agroinfiltration. The samples were harvested 3 hours after biotin and estradiol treatment.

#### Measurement of electrolyte leakage

For measurement of ion leakage, Arabidopsis leaf discs (6-8 mm diameter) were collected 48 hours post dex treatment and were immersed in a 50 ml sterile tube containing 15ml of water. For transient assays in *N. benthamiana*, leaf discs (9-10mm) collected from the agro-infiltrated region at either 48 or 72 HPI were immersed in a 50 ml sterile tube containing 15ml of water. Conductance was measured at the indicated time points using a conductivity meter (WTW, Weilheim, Germany). In both cases, leaf discs from non-infiltrated leaves were also collected to establish background ion leakage. Each assay included 3-5 separate biological replicates, with 2-3 technical replicates measured at a given time point. For data presented as percent ion leakage, samples were measured then boiled and remeasured to determine maximum ion leakage.

### Confocal microscopy

*N. benthaminana* leaves were infiltrated with Agro carrying the indicated construct(s). At 48 or 72 hrs after infiltration, YFP and/or RFP florescence were observed with a confocal microscope (Nikon Eclipse Ti/C2/C2Si) (Nikon, Foster City, CA) using an excitation wavelength of 514 nm and emission wavelength of 530 nm or an excitation wavelength of 561 nm and an emission wavelength of 575 nm, respectively. Nuclei were stained by infiltrating 0.4ug/ml Hoechst 33342 dye (Sigma-Aldrich, St. Louis, MO) into leaves 8 hours before image acquisition ^76^ and observed using an excitation wavelength of 350 nm and an emission wavelength of 460 nm.

### Trypan Blue Staining

Trypan blue staining was performed as described ^27^. Briefly, staining solution was prepared by mixing 1 part staining mix (1:1:1:1 mix of phenol, lactic acid, glycerin, and water plus 0.05% [w/v] trypan blue) and 2 parts ethanol. The leaves were submerged in the staining solution for 3 min at 95°C followed by an additional overnight incubation. Stain was removed by 15 M chloral hydrate solution and mounted in 70% glycerol.

## Resource availability

### Lead contact

Further information and requests for material should be directed to the lead contact: David Mackey

## Acknowledgements

35S:RFP-OsRac1 was a gift from Dr Guo-Liang Wang. HA-NDR1 was a gift from Dr Brad Day. 35S:AvrRpt2-HA and 35S:AvrRpt2^C122A^-HA were a gift from Dr Kee Hoon Sohn. The Mackey laboratory is supported by the National Science Foundation (Division of Integrative Organismal Systems, grant no. 1953509). J.H. was supported by the EUs Horizon 2020 research and innovation program under the Marie Skłodowska-Curie scheme (No 897584). The Jones lab is supported by the Gatsby Charitable Foundation (Core grant to TSL).

## Author contributions

D.M., A.J.A and L.C. conceived the project. D.M. and A.J.A supervised the experiments. A.J.A, M.A. and J.H. designed and performed the experiments. Additionally, L.C., M.A.A, M.I.R, J.T. and A.J. contributed towards conducting experiments. A.J.A, M.A., J.H. and M.A.A. analyzed the data. Resources for the experiment were provided by A.J.A, J.D.G.J and D.M. D.M., A.J.A. and M.A.A wrote the original draft. The final version was reviewed and edited by M.A., J.D.G.J. and D.M.

## Supplemental information

Document S1. Figure S1-10

