## Supplement figures for "Defense-Suppressive Fragments of RIN4 generated by AvrRpt2 Participate in NDR1-dependent Activation of RPS2"

**Supplemental Figures:**

**Figure S1:** Membrane-tethered derivatives of RIN4 that contain an NOI domain cause chlorosis, but not cell death. **A)** Expression of RIN4 derivatives in homozygous transgenic Arabidopsis plants was induced by dex. Macroscopic symptoms were observed at 72 HPI. **B)** Trypan blue staining of leaves from plants as in **A**. **C)** Cell death in plants as in **B** was quantified by measuring electrolyte leakage. Data was collected from 4 independent experiments with at least 2-3 technical replicates per transgenic line (n=3-12). Error bars represent SEM. **D)** Anti-T7 western blot shows accumulation of RIN4 derivatives at 72 HPI in the indicated transgenic lines. The lower panel shows Coommasie brilliant blue stain for RuBisCO used as loading control. Asterisks indicate the positions of individual RIN4 derivatives.

**Figure S2**: Suppression of RPS2 in *N. benthamiana* by membrane-tethered derivatives of RIN4 depends on C-NOI and, to a lesser extent, N-NOI. **A)** Flag-RIN4 derivatives expressed from 35S promoter in transient experiments in *N. benthamiana*. **B)** The indicated derivatives of RIN4 ($OD_{600}$ = 1.0) and RPS2-HA ($OD_{600}$ = 0.05) were co-infiltrated in *N. benthamiana*. Macroscopic symptoms were observed at 48 HAI. **C)** Cell death in plants as in **B** was quantified by measuring electrolyte leakage at 96 HAI. Data was collected from 3 independent experiments with 3 technical replicates per treatment (n=9). Error bars represent SEM. Student's t-test, at 95% confidence limits, was used for comparison with Flag-RIN4Fl (ns, not significant; *P<0.05, **P<0.01; ***P<0.001). **D)** Anti-Flag immunoblot shows accumulation of Flag-RIN4 derivatives expressed in *N. benthamiana* at 72 HAI. The lower panel shows ponceau stain for RuBisCO used as loading control. Asterisks indicate the positions of individual RIN4 derivatives.

**Figure S3:** Non-membrane-tethered derivatives of RIN4 depend on their C-NOI domain to cause a cell death response in Col-0. **A)** Expression of RIN4 derivatives was induced by dex treatment in homozygous transgenic Arabidopsis plants. Macroscopic symptoms were observed at 72 HPI. **B)** Trypan blue staining of leaves from plants as in **A**. **C)** Cell death in plants as in **A** was quantified by measuring electrolyte leakage. Data was collected from 4 independent experiments with at least 3 technical replicates per transgenic line (n=6-12). Error bars represent SEM. **D)** Anti-T7 immunoblot showing protein accumulation of RIN4 derivatives in the indicated transgenic lines at 72 HPI. The lower panel shows coommasie brilliant blue stain of RuBisCO as a loading control. Asterisks indicate the positions of individual RIN4 derivatives.

**Figure S4:** Overexpression of AtNDR1 in *N. benthamiana* suppresses RPS2-induced cell death. **A)** Flag-RIN4Fl, RPS2-HA, HA-NDR1 and AvRpt2-HA constructs used in transient experiments in *N. benthamiana*. **B)** The indicated combinations of Flag-RIN4Fl ($OD_{600}$ 0.6), HA-NDR1 ($OD_{600}$ = 0.6), AvrRpt2-HA and/or AvrC122A-HA ($OD_{600}$ = 0.01) and RPS2-HA ($OD_{600}$ = 0.08), were co-infiltrated in *N. benthamiana*. Macroscopic symptoms were observed at 48 HAI. **C)** Cell death in plants as in **B** was quantified by measuring electrolyte leakage at 72 HAI. Error bars represent SEM. Data was collected from 3 independent experiments with 3 technical replicates per treatment (n=9). Student's t-test, at 95% confidence limits, was used for comparison with Flag-RIN4Fl (ns, not significant; *P<0.05; **P<0.01; ***P<0.001). **D)** The indicated combinations of RIN4 derivatives ($OD_{600}$ 0.6), with HA-NDR1 ($OD_{600}$ = 0.6) and RPS2-HA ($OD_{600}$ = 0.05), were co-infiltrated in *N. benthamiana*. Macroscopic symptoms were observed at 48 HAI. **E)** Cell death in plants as in **D** was quantified by measuring electrolyte leakage at 96 HAI. Error bars represent SEM. Data was collected from 4 independent experiments with 3 technical replicates per treatment (n=12). Student's t-test, at 95% confidence limits, was used for comparison with Flag-RIN4Fl (ns, not significant; *P<0.05; **P<0.01).

**Figure S5**. Activation of RPS2 in *N. benthamiana* by non-membrane-tethered RIN4 requires its accumulation in the cytosol. **A)** YFP taggedRIN4 derivatives expressed from 35S promoter in transient experiments in *N. benthamiana*. N-terminal rectangles, NLS (green), NES (grey), or SNE (orange) tags. **B)** The indicated RIN4 derivatives were expressed in *N. benthamiana*. Shown are YFP, Hoechst, and merged confocal microscope images at 48 HAI and following Hoechst staining. **C)** Anti-GFP immunoblot shows accumulation of RIN4 derivatives expressed in *N. benthamiana*. The lower panel shows ponceau staining for RuBisCO, used as loading control. Asterisks indicate the positions of individual RIN4 derivatives. **D)** The indicated RIN4 derivatives or free YFP were co-expressed with RFP-OsRac1 in *N. benthamiana*. Shown are YFP, RFP, and merged confocal microscope images at 72 HPI. Scale bars are 15 μM. **E)** The indicated combinations of Flag-RIN4Fl ($OD_{600}$ = 1.0), either YFP-NLS-CCC>AAA ($OD_{600}$ = 0.6 or 0.3) or YFP-NES-CCC>AAA ($OD_{600}$ = 0.3) and RPS2-HA ($OD_{600}$ = 0.07), were co-infiltrated in *N. benthamiana*. Macroscopic symptoms observed at 48 HAI. **(F-G)** Cell death in plants as in **E** was quantified by measuring electrolyte leakage at 48 HAI. Error bars represent SEM. Student's t-test, at 95% confidence limits, was used for comparison with Flag-RIN4Fl (**Right panel-**ns, not significant; **P<0.005; ***P<0.001).

(**Left panel-**ns, not significant; **P<0.01; ***P<0.005). (H) Anti-YFP western blot shows expression levels of YFP-NLS-CCC>AAA and YFP-NES-CCC>AAA at 48 HPI. Panels below show ponceau stain for RuBisCO used as loading control. Arrow indicates positions of the RIN4 derivatives.

**Figure S6**: RPS2-TurboID-V5 specifically biotinylates Flag-RIN4 and Ac5V-NES-RIN4. **A)** tagged 35S:RPS2-Turbo-ID-V5 and 35S:BRI1-Turbo-ID-V5 constructs. **B)** RIN4 derivatives and either RPS2-TurboID-AcV5 or Bri1-TurboID-V5 (each at $OD_{600}$ = 0.5) were co-infiltrated in *N. benthamiana* leaves. After 24hrs the leaves were infiltrated with biotin and 4hrs later the tissue was collected. A portion was immunoprecipitated with anti-strep and the input and IP samples were immunoblotted as indicated with red arrows indicating the position of the proteins. **C)** RIN4Fl, NES-RIN4-CCC>>AAA and either RPS2-TurboID-V5 or Bri1-TurboID-V5 were co-infiltrated in *N. benthamiana* leaves (each at $OD_{600}$ = 0.5) and were processed as in A. V5-HRP was used to detect RPS2 and BRI1, while Flag-HRP and anti-AcV5 was used for detection of biotinylated proteins (bottom panels).

**Figure S7.** ACP2 is unable to activate RPS2 suppressed by full-length RIN4 or ACP3 with a bulky N-terminal tag in *N. benthamiana.* **A)** The indicated RIN4 derivatives ($OD_{600}$ = 0.6) and RPS2-HA ($OD_{600}$ = 0.04) were co-infiltrated in *N. benthamiana*. Macroscopic symptoms were observed at 48 HAI. **B)** Cell death in plants as in **A** was quantified based on electrolyte leakage. Leaf discs were immersed in sterile water and conductivity was measured at 96 HAI. Data was collected from 6 independent experiments with 3 technical replicates per treatment (n=18). Error bars represent SEM. Student's t-test, at 95% confidence limits, was used for comparison with Flag-RIN4Fl (ns, not significant; **P>0.01). **C)** The indicated RIN4 derivatives ($OD_{600}$ = 0.6) and RPS2-HA ($OD_{600}$ = 0.04) were co-infiltrated in *N. benthamiana*. Macroscopic symptoms were obaerved at 48 HAI. **D)** Cell death in plants as in **C** was quantified by measuring electrolyte leakage 96 HAI. Data was collected from 3 independent experiments with 3 technical replicates per treatment (n=9). Error bars represent SEM. Student's t-test, at 95% confidence limits, was used for comparison with Flag-RIN4Fl (ns, not significant; **P<0.01). **E)** RIN4 derivatives and either RPS2-TurboID-AcV5 or Bri1-TurboID-AcV5 (each at $OD_{600}$ = 0.5) were co-infiltrated in *N. benthamiana* leaves. After 24hrs the leaves were infiltrated with estradiol. 8hrs later, the same leaf was infiltrated with biotin and 4hrs post biotin infiltration the tissue was collected. A portion was immunoprecipitated with anti-strep and input and IP samples were immunoblotted as indicated. Red arrows indicate the position of the proteins. Anti-V5-HRP was used to detect RPS2 and BRI1, while anti-GFP-HRP and Flag-HRP were used for detection of biotinylated proteins.

**Figure S8:** Schematic diagram of homologous RIN4 derivatives. Within RIN4 derivatives: Yellow rectangles indicate YFP tag, peach rectangle indicates AcV5 epitopes, pink triangle indicates Flag epitope, purple rectangles indicate NOI domains, and blue triangles indicate RCS.

**Figure S9:** Spatial proximity of RPS2 and ADR1-L1 in *N. benthamiana.* ADR1-L1-Flag and either RPS2-TurboID-V5 or Bri1-TurboID-V5 (each at $OD_{600}$ = 0.5) were co-infiltrated in *N. benthamiana* leaves. After 24hrs the leaves were infiltrated with estradiol. 8hrs later, the same leaf was infiltrated with biotin and 4hrs post biotin infiltration the tissue was collected. A portion was immunoprecipitated with anti-strep and input and IP samples were immunoblotted as indicated. Red arrows indicate the position of the proteins. **A)** Anti-V5-HRP was used to detect RPS2 and BRI1, while anti-Flag-HRP was used for detection of ADR1-L1. **B)** Streptavidin HRP was used to detect biotinylated proteins.

**Figure S10:** The schematic diagram of AvrRpt2 induced activation of RPS2 in *Arabidopsis*. In the absence of the pathogen, membrane localized RIN4 suppresses RPS2 activation. *P. syringae* secretes AvrRpt2, a cysteine protease, in plant cytosol using the T3SS. AvrRpt2 cleaves RIN4 at the conserved motif VPXFGXW, known as RCS1 and RCS2 (RIN4 cleavage sites), in the N and the C-NOI domains respectively. Cleavage of RIN4 at these sites results in the generation of three fragments, namely ACP1, ACP2 and ACP3. The production of non-membrane tethered fragment, ACP2, elicits RPS2 activation in the presence of the membrane tethered fragment, ACP3. Activated RPS2 in turn triggers defense gene expression and HR.
