## Supplementary figures and images for "Defense-Suppressive Fragments of RIN4 generated by AvrRpt2 Participate in NDR1-dependent Activation of RPS2"

### Figure S1

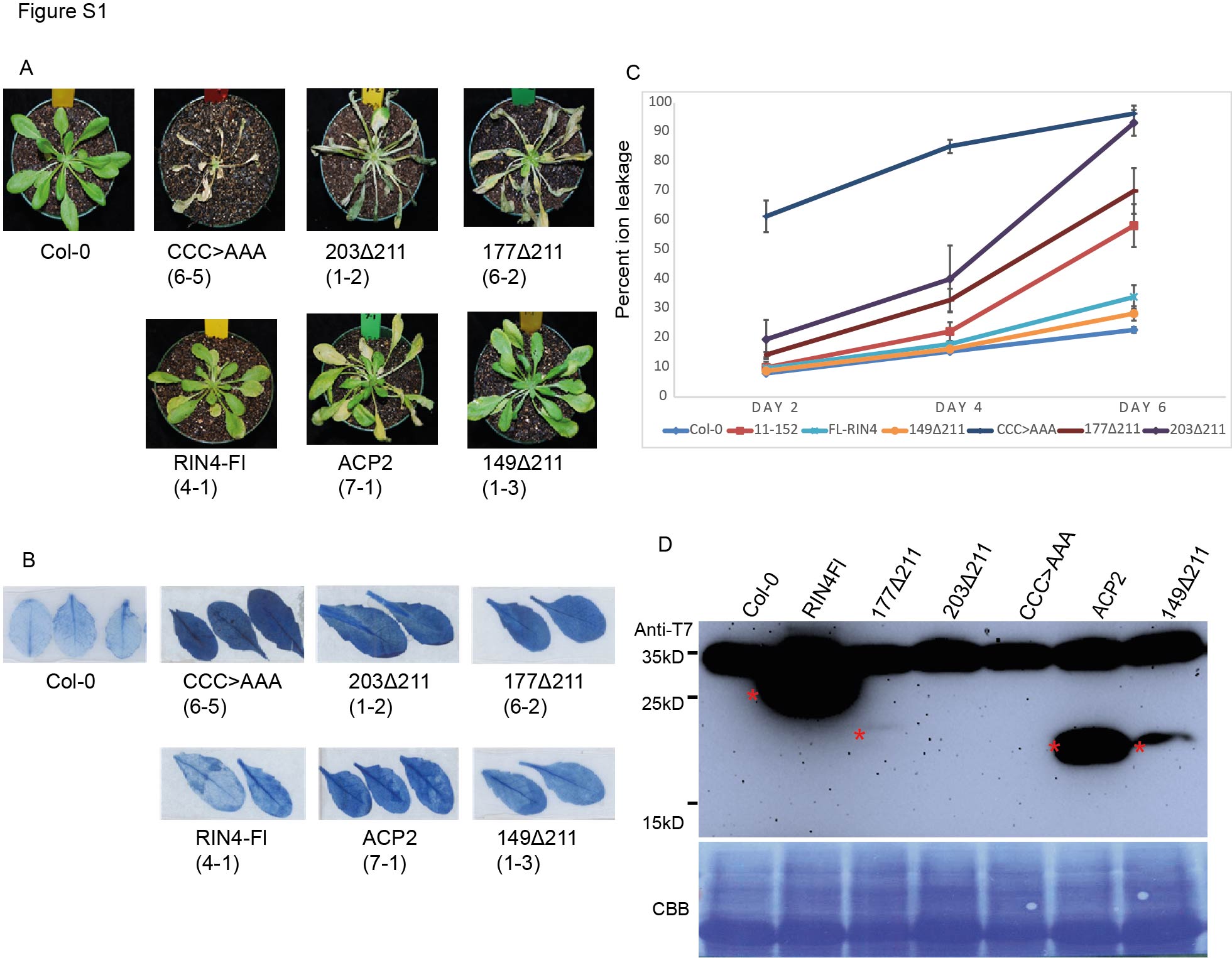

### Figure S2

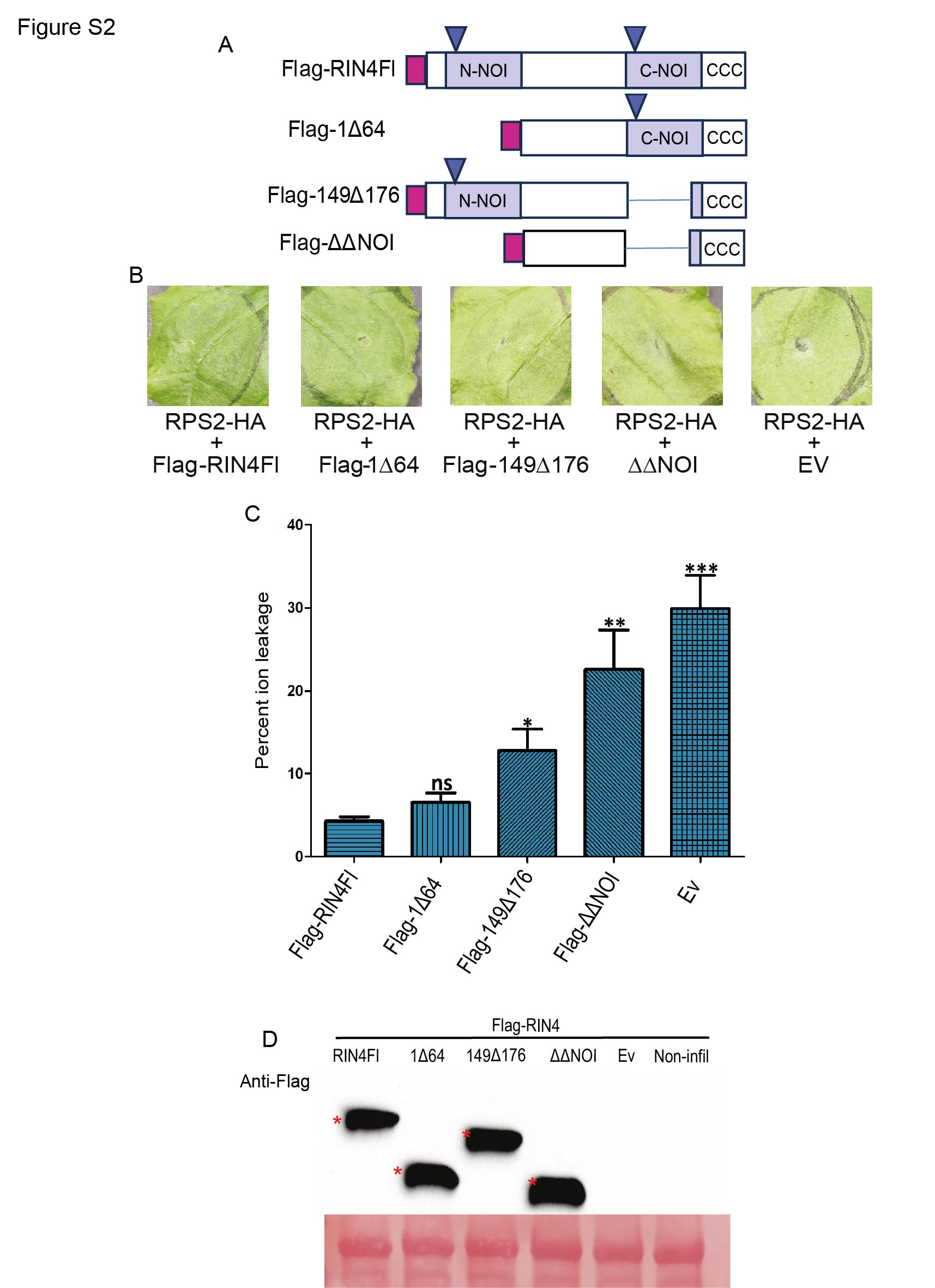

### Figure S3

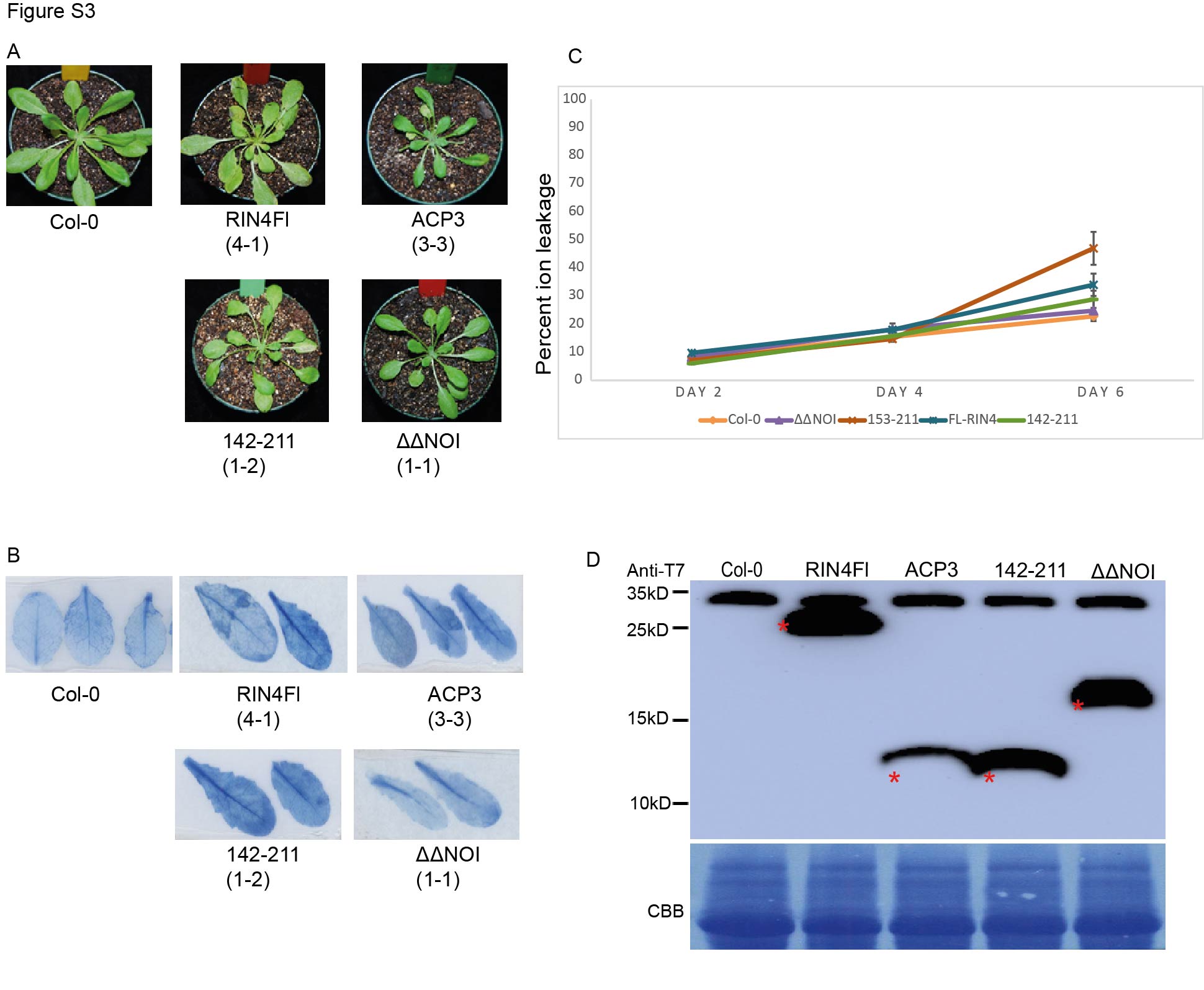

### Figure S4

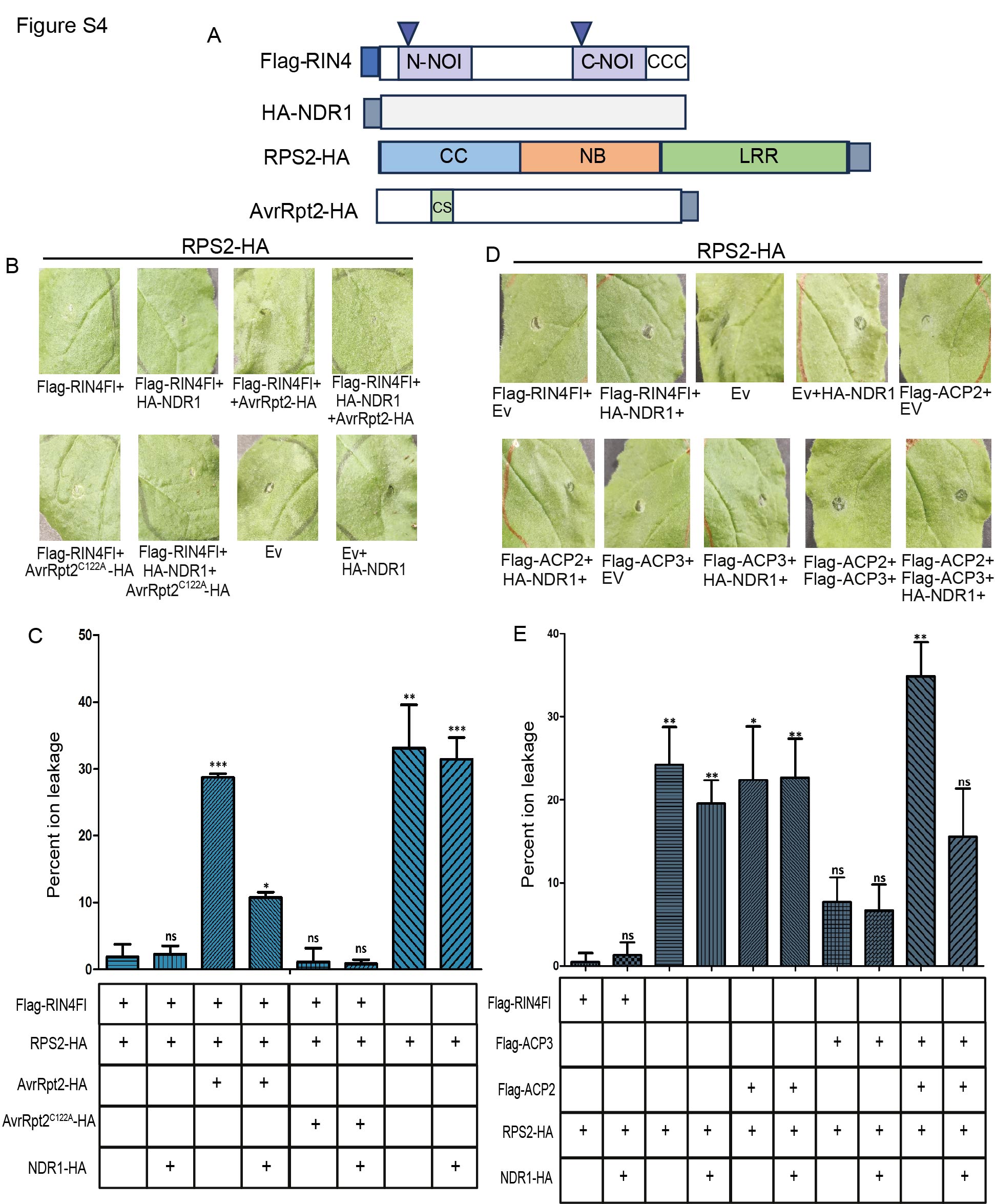

### Figure S5

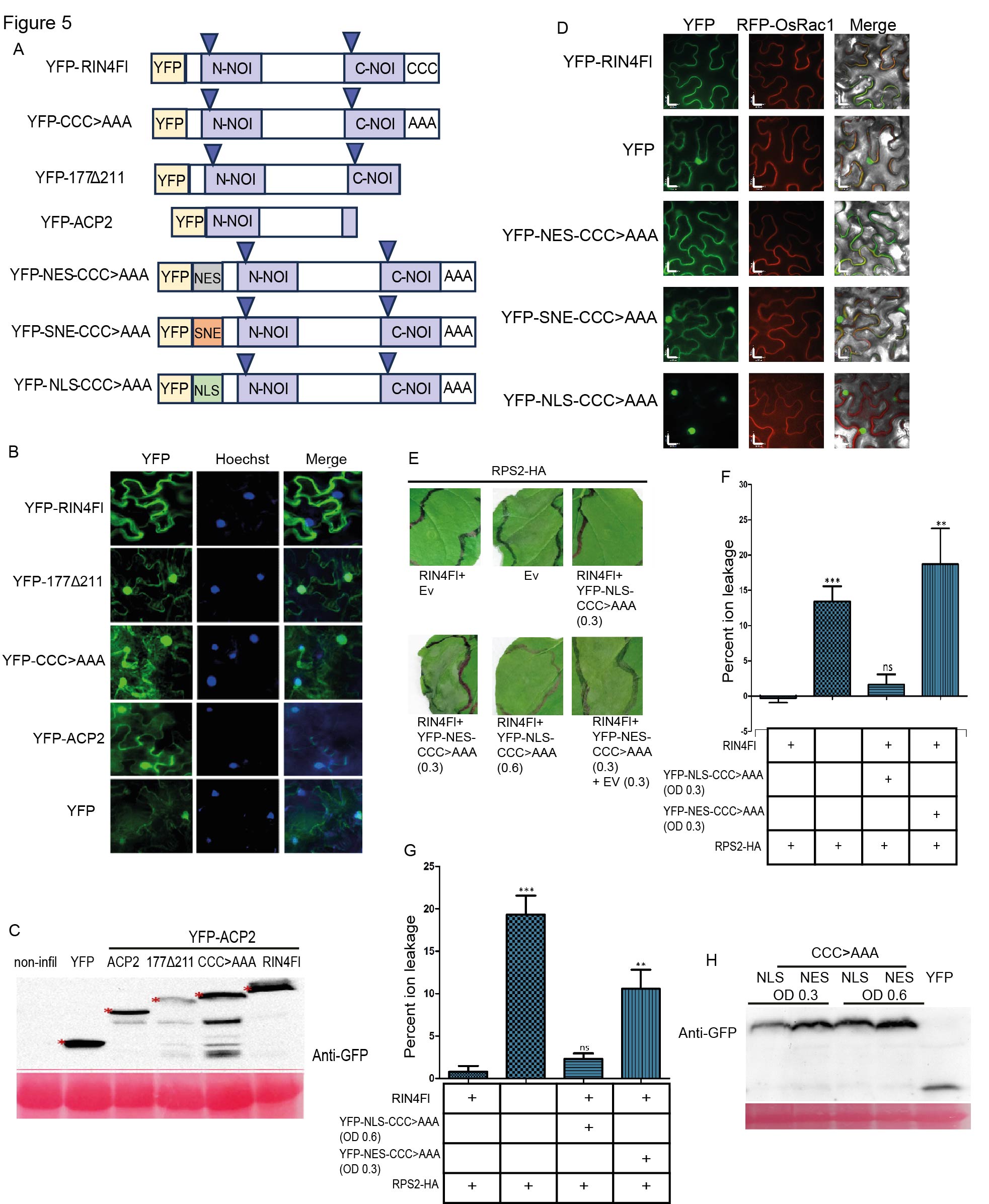

### Figure S6

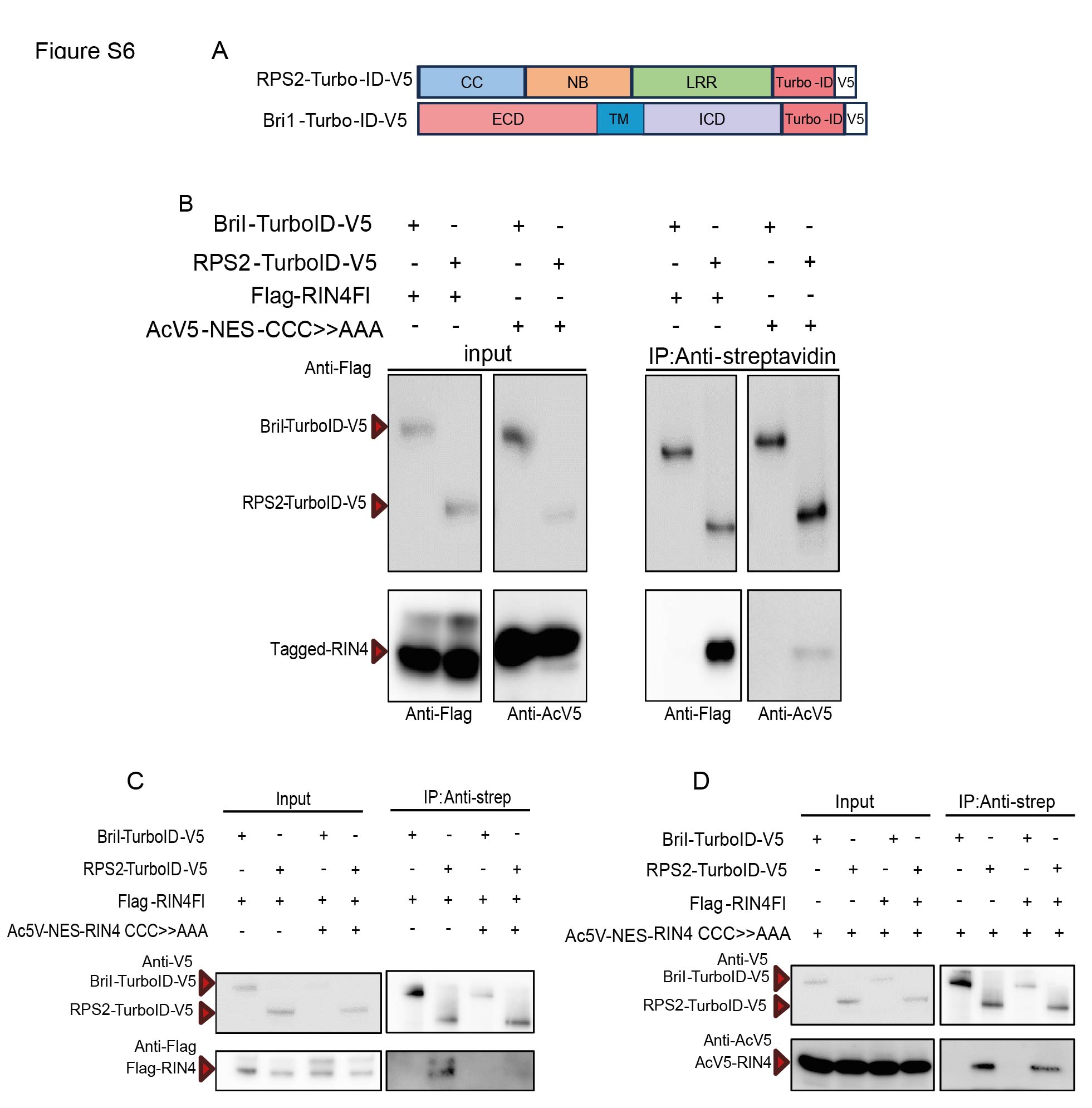

### Figure S7

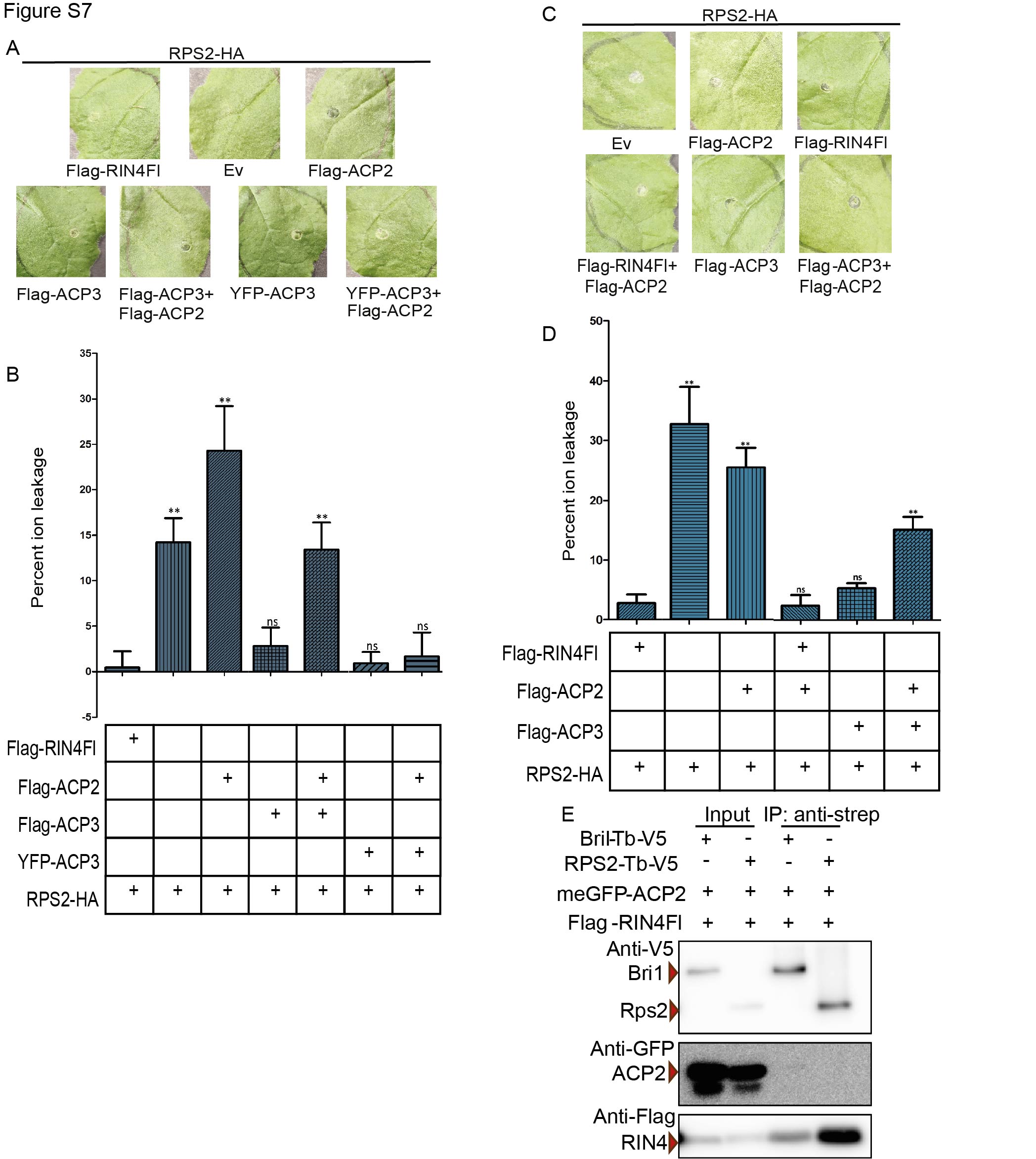

### Figure S8

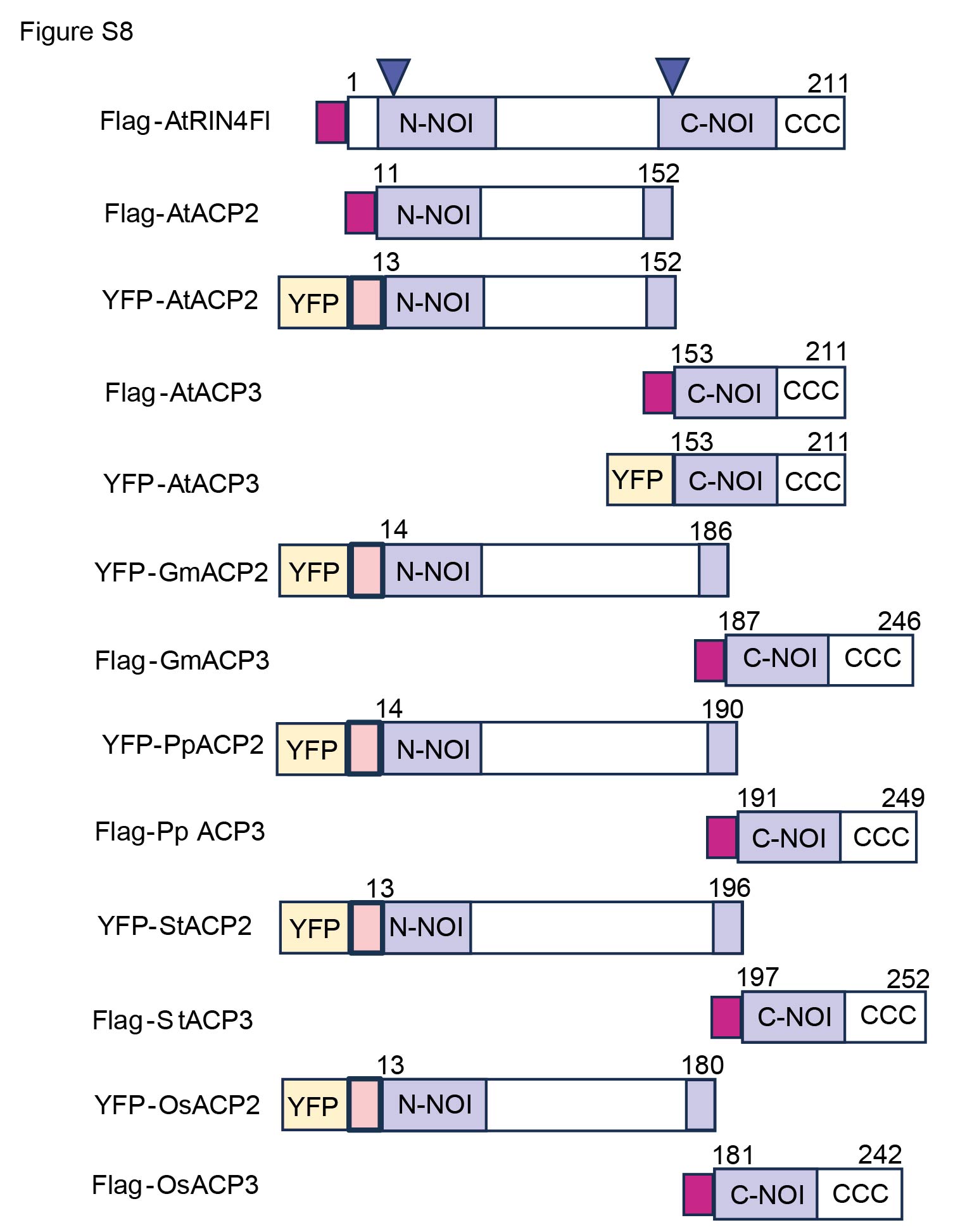

### Figure S9

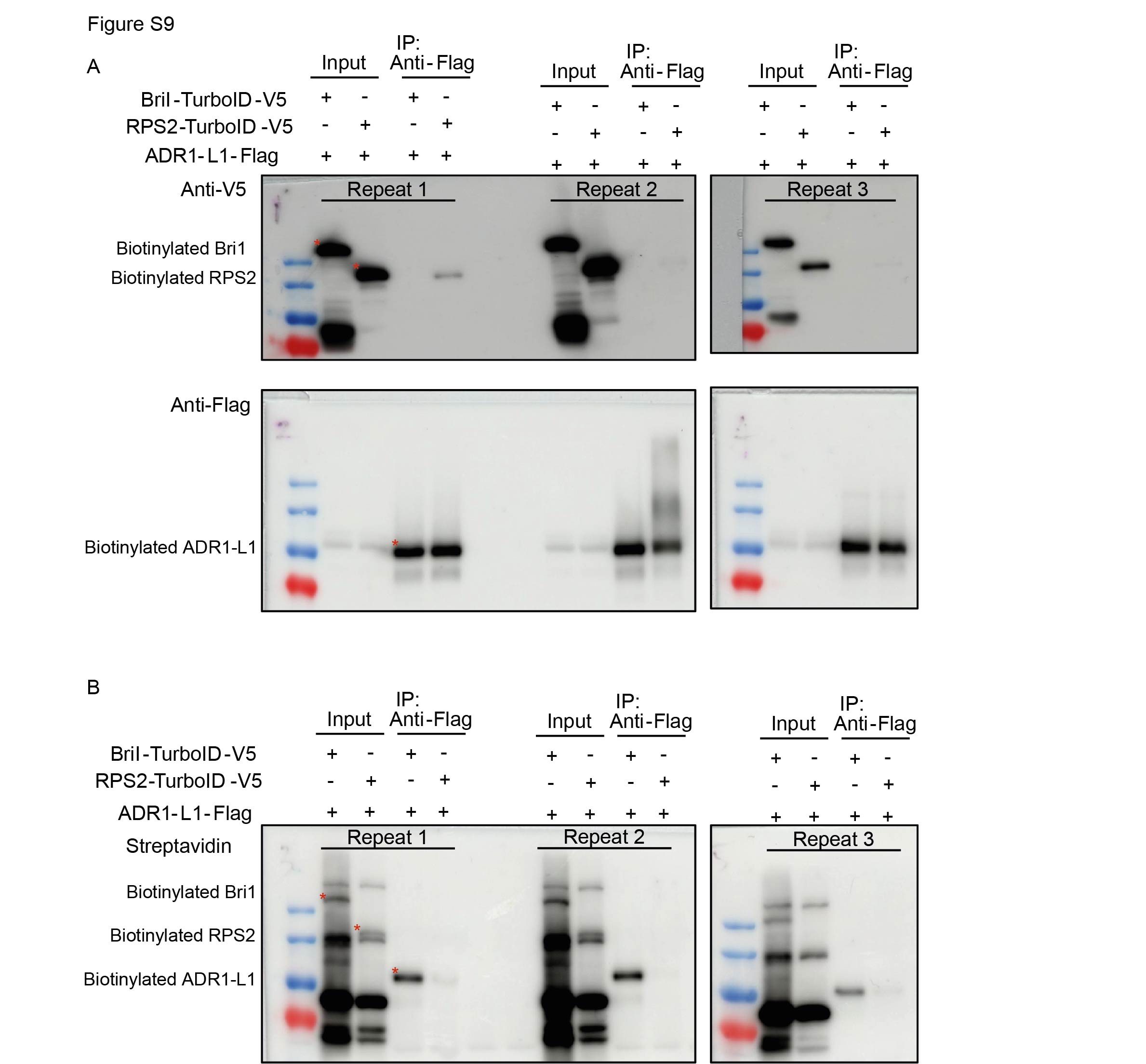

### Figure S10

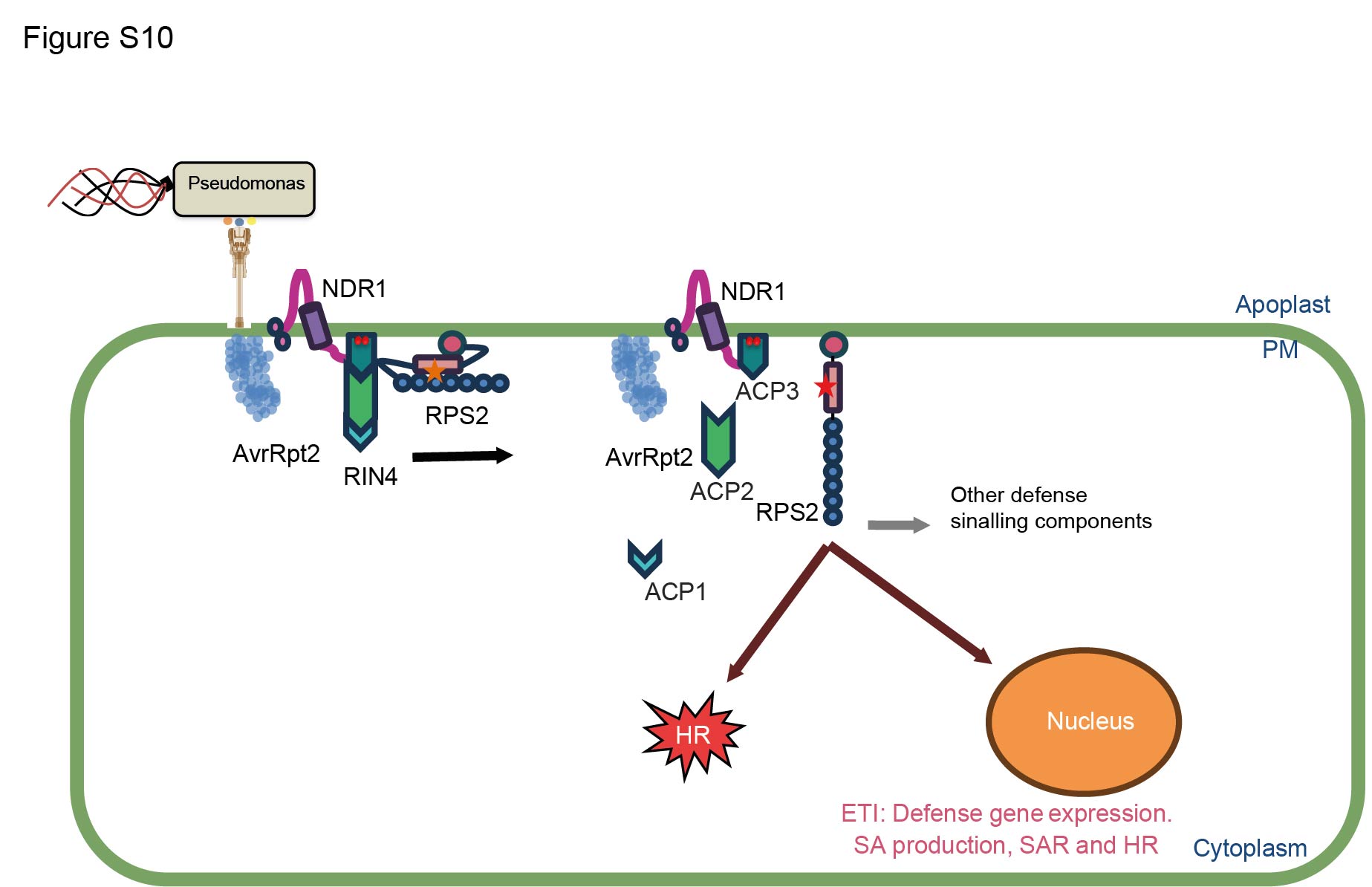
